# Pathways to cultural adaptation: the coevolution of cumulative culture and social networks

**DOI:** 10.1101/2023.02.21.529416

**Authors:** Marco Smolla, Erol Akçay

## Abstract

Humans have adapted to an immense array of ecologies by accumulating culturally transmitted knowledge and skills. Culture accumulates in at least two ways: via more distinct cultural traits, or via improvements of existing cultural trait. A trade-off is expected between these owing to the fact that social learning opportunities are finite and social learning often requires multiple exposures. Furthermore, what kind of culture accumulates depends on, and coevolves with, the social structure of societies. Here we show that the coevolution of social networks for learning and cumulative culture results in two distinct pathways to cultural adaptation: highly connected populations with high proficiency but low cultural trait diversity vs. sparsely connected populations with low proficiency but more cultural trait diversity. Importantly, we show there is a general conflict between group-level payoffs, which is maximised in highly connected groups that attain high proficiency, and individual level selection, which favours disconnection. This conflict emerges from the interaction of social learning with population structure and causes populations to cycle between the two cultural and network states. The same conflict creates a paradox where improving individual innovation rates lowers the payoffs of groups. Finally, we explore how populations navigate these two pathways in heterogeneous and changing environments, and show that high heterogeneity in payoffs and slow rate of environmental change favours high proficiency, while fast rate of environmental change favours more trait diversity. We also find that the proficiency pathway to cultural adaptation is favoured with increased population size, but only in slow changing environments. Our results uncover previously unrecognised trade-offs and tensions in the coevolutionary dynamics of cumulative culture and social structure, with broad implications for human social evolution.

## 1 Introduction

Our species’ ecological success is in large part based on our capacity to accumulate cultural knowledge (Boyd et al., 2011; Henrich and McElreath, 2003; Hill et al., 2009). We have accumulated vast amounts of cultural traits (e.g., knowledge, technologies, believes, and ideas) that are too numerous or complex to be invented by a single individual (Boyd and Richerson, 1995; Henrich and McElreath, 2003; Mesoudi and Thornton, 2018). Instead, cumulative cultural adaptation proceeds by individual innovations that either add a new skill, tool, or knowledge, or add to the complexity and efficacy of an existing one. These individual innovations, if they spread to enough individuals through social transmission, can be maintained in the population in the long term so that they do not have to be reinvented and can be built upon. In fact, several researchers have pointed at our specie’s unique cultural niche, which is build on various forms of social learning that allow us to specialise on resources that require high skill, and do so in a broad spectrum of environments (Hill et al., 2009; Kaplan et al., 2000; Roberts and Stewart, 2018). This capacity—to accumulate locally adaptive traits—enabled us to settle all over the globe from arid deserts to frigid polar regions, and from the rich equator to the relatively unproductive high-latitudes (Boyd et al., 2011; Elton, 2008).

Whether and how much adaptive culture accumulates is a function of social and demographic parameters, such as group size (Derex et al., 2013; Henrich, 2004b), connectivity (Cantor et al., 2021; Derex et al., 2018), life span (Acerbi et al., 2012), acquisition costs (Mesoudi, 2011), and length of the learning period (Lehmann et al., 2010). The converse is also true: the quantity and quality of cumulative culture and the selection on cultural traits will determine selection on individual-level and group-level traits such as learning schedules (Lehmann et al., 2013) or network structure (Smolla and Akçay, 2019). Thus, understanding adaptation through cumulative culture needs to consider the reciprocal feedbacks between the dynamics of innovation and transmission of cultural traits and the individual and group-level mechanisms through which these dynamics occur.

An important feature of cumulative culture is that there are often many cultural traits that might be profitable, and each cultural trait can be built upon and improved by successive innovations. This gives individuals and societies multiple routes to increase their payoffs: (1) innovating and/or learning more profitable traits (i.e., having a large repertoire) or (2) innovating and/or learning improvements on traits that they already have (i.e., having high proficiency in fewer traits). Our previous work showed that when cultural selection favours large repertoires, groups evolve sparsely connected networks and large trait diversity, whereas selection for high proficiency results in densely connected groups that coordinate on a few traits, allowing successive innovations to be maintained in the group (Smolla and Akçay, 2019).

In the real world, groups and individuals can benefit from either accumulating more traits or higher proficiency—or both. Such open-ended cultural selection creates an inherent tradeoff between learning new traits versus improving proficiency on existing ones. As cultural traits and proficiency accumulates in the population, individuals will be constrained by limits of social learning, especially when social learning requires multiple exposures to the trait to be learned. The consequences of this inherent trade-off has not yet been explored. Most studies of cumulative culture rely on either accumulation of a single dimension of cultural complexity (e.g. Henrich, 2004a), with only a few studies (Kempe et al., 2014; Kolodny et al., 2015a; Mesoudi, 2011) that consider the accumulation and improvement of different cultural traits. In most of these studies however, there is no trade-off between learning new traits or maintaining and improving existing ones (Mesoudi, 2011, is a notable exception). Moreover, most models of cumulative culture regard social learning as a simple contagion process, assuming spread proportional to their local prevalence. This corresponds to an effective assumption that social learning happens or not instantaneously upon exposure. However, traits that make up adaptive cumulative culture, such as foraging tactics or making tools, are likely to require repeated exposures to be learned. This makes spread of cultural traits a non-linear function of their local prevalence. Our own previous work (Smolla and Akçay, 2019) incorporated this non-linearity and a trade-off at the individual level but sidestepped it at the population level by exogenous imposing selection for either only broad repertoire or only high proficiency. As a result, how individuals and societies navigate the competing pathways to cultural adaptation, and how these pathways coevolve with social structure remains largely unknown. As we show here, this coevolution is subject to an unexpected emergent conflict between individual and group level cultural adaptation.

Another fundamental feature of cumulative cultural evolution is that it almost always happens in heterogeneous environments where the payoffs from different traits will be variable and fluctuate over time. Thus, cultural traits that are profitable in a given time and place might not be profitable in another (Boyd and Richerson, 1995; Richerson and Boyd, 2005). Such fluctuations can alter the trade-off between broader repertoires and higher proficiency: environmental change is expected to favour broader, more generalist cumulative culture as a way of bet-hedging (Deffner and Kandler, 2019) which in turn might be expected to favour individual innovation and cultural diversity. However, how cumulative culture coevolves with social network structure in heterogeneous and fluctuating environments remains unknown.

To address these gaps, we model open-ended cultural selection where individuals can increase their payoffs either by learning new cultural traits or increasing their proficiency in traits they already know. We first ask what kinds of social structure is produced by such open-ended cultural selection and find that counter-intuitively, societies tend to cycle between densely connected states with high-proficiency culture and sparsely connected states with broad-repertoire culture. Interestingly, this cycling happens despite the fact that the specialist state has a much higher overall payoff. We show that cycling is driven by a conflict between group-level cultural adaptation and individual selection: average group level payoff is highest in highly connected, high proficiency culture, but individual level selection favours sparser connections that eventually break apart the high-proficiency culture. Likewise, we show that increasing individual innovation rate inhibits high-proficiency culture through the same mechanism. We then ask what kind of social network structure and cumulative culture coevolve in heterogeneous and fluctuating environments where cultural traits have variable payoffs that change over time. We find that higher rates of environmental change and lower variation in payoffs between traits favour generalist societies with sparse network connections, while slower environmental change and higher trait variance favour specialist and well-connected societies. These results illustrate the complex interplay between group structure, social learning, and environmental variation in cumulative cultural adaptation.

## 2 Methods

We consider a population of *N* asexually reproducing individuals, with overlapping generations in a world with *T* learnable traits or skills. These skills relate to subsistence, social norms or other aspects relevant to an individual’s survival. As we are interested in the effect of environmental heterogeneity on social structure and cultural evolution, we assume that skills differ in their utility. A skill’s utility, *u*, translates into a payoff that an individual receives if the skill is part of its repertoire. The values of *u* are randomly drawn for each trait, *t*, from lognormal distributions, Lognormal(*μ, σ*^2^) in the interval [0,10]. If not stated otherwise we use *μ* = 0 and *σ*^2^ ∈ {0.2,0.4,..., 1}, and so mainly varying payoff variance (see supplementary material, Fig. S1). We simulate environmental change as a change in skill utilities across time. In our model, time is divided into rounds. In each round each skill has a small probability to receive a new randomly drawn utility from the same distribution the simulation was started with. We express the environmental change probability as the number of updated utilities per generation (i.e., every N rounds) *τ* ∈ {10^-3^,10^-2^,..., 10^1^}. In fast changing environments (*τ* = 10^1^), 10 utilities are updated every generation. At the other extreme (*τ* = 10^-3^) it takes on average 1000 generations for a single utility to change.

In addition to environmental change, in each round: (1) an individual is removed from the population (selected relative to the inverse of their payoff), (2) a parent is randomly selected and a copy (subject to mutation of the connection traits) is added to the social network, and (3) the new individual acquires cultural traits and proficiency through innovation (individual learning) or social learning.

### 2.1 Population structure

We use complex dynamic networks structured by social inheritance (Ilany and Akcay, 2016) to simulate population structure and turnover. These networks capture important aspects of real-world networks (Ilany and Akcay, 2016) and allow the dynamical formation of local and global clustering in response to different selective regimes. The social inheritance model has three linking parameters, which represent the probabilities that a new individual (1) forms a connection with its parent, *p*_b_ (here, we assume *p*_b_ **= 1**), (2) with the neighbours of the parent, *p*_n_, and (3) with other individuals that are not connected to the parent, *p*_r_. A new individual that is added to the population inherits *p*_n_ and *p*_r_ vertically from its parent (culturally or genetically see, e.g., Brent et al., 2013). Mutation occurs with probability *m* = 0.05, whereby mutated values are drawn from a normal distribution centred around the parent’s value with standard deviations 0.05 and 0.005 for *p*_n_ and *p*_r_ respectively. The new individual is then connected to its parent and to other individuals based on *p*_n_ and *p*_r_.

### 2.2 Learning

Next, the new individual enters the learning phase, allowing her to either acquire new skills or improve skill proficiency. The learning phase consists of 100 social learning and 100 innovation attempts. At birth, an individual’s proficiency *l* is zero for all *T* skills. Skill proficiency increases through successful learning. When a new skill is acquired through innovation or copying proficiency of skill *t* increases by one unit, i.e. *l_t_*\* = *l_t_* + 1. An individual’s repertoire size *R_i_* is the number of non-zero skill proficiencies *l*. To become better at performing a skill, repeated engagement with it is required, as learning takes time (Karni et al., 1998; Karni and Sagi, 1993; Lew-Levy et al., 2017; Morelli et al., 2003). Therefore, proficiency increases with each successful individual or social learning attempt of the same skill. An individual’s highest proficiency *L_i_* is equal to the largest *l_t_* in its repertoire. Because the number of learning turns is limited and attention to one skill limits attention to other skills, there is a trade-off between becoming good at a skill and learning many skills. Hence, there is a negative relationship between skill proficiency and repertoire size.

During an *individual learning* episode the individual first picks one skill from all possible skills *T* at random, and then attempts to acquire proficiency for this skill. Learning success is moderated by an innovation success probability, *α*, and so the probability to acquire *t* through innovation is

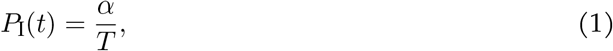

During a *social learning* episode the individual first picks a skill from those performed in its vicinity (i.e. present in the repertoire of her neighbours). Given that each individual is assumed to be equally likely to perform any of their traits, the probability that individual i observes trait t in her neighbourhood is 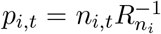, where *n_i,t_* is the number of *i*’s neighbours with skill *t*, and *R_n_i__* is the sum of repertoire sizes of *i*’s neighbours. Subsequently, the newborn attempts to acquire proficiency for this skill. Learning success is moderated by a social learning success probability, *β*. Crucially, we assume that social learning is more effective when an individual receives more exposure to a skill, i.e., if *p_i,t_* is larger, and so the probability to acquire t through copying is

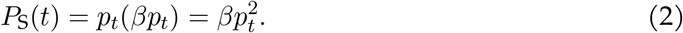

Equation 2 can be thought of as complex contagion in a given trait (Centola, 2010), where transmission depends on the probability of observing a skill twice, regardless of the individual being observed. As pointed out previously (Smolla and Akçay, 2019), this relates to Simpson’s index (Simpson, 1949), and so as skill diversity in *i*’s neighbourhood increases the exposure to each skill decreases, making it less likely to be observed sufficiently for successful social learning. Thus, equation 2 shows acquiring a skill socially is more likely if social learning is easy (large *β*), *t* is common among neighbours (large *n_i,t_*), and/or if neighbours possess few skills (small *R_n,i_*). Additionally, we assume that an individual cannot surpass the proficiency of the observed individuals by social learning, and thus *P*_S_(*t*) = 0 where all neighbours have proficiency equal to or less than that of the individual *i* for skill *t*.

### 2.3 Payoff

After the learning phase, we calculate the total payoff *W*, which an individual receives from each skill in its repertoire. We calculate *W_i_* as the sum of the product of the skill proficiencies in *i*’s repertoire and their utility:

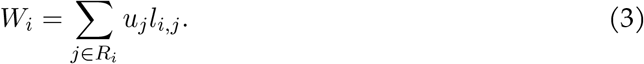

Because of the trade-off between learning many different skills or becoming very good in a few skills, Eq. 3 can be maximised in two different ways: either by acquiring some proficiency in many well paying traits (hereafter *generalist*), or by becoming proficient in the most profitable skills (hereafter *specialist*). A homogeneous environment (where *u_j_* is the same for all *j*), learning a new trait or improving proficiency yields the same payoff, so our utility function does not inherently favour one or the other pathway.

We model here mortality selection due to cultural traits, where *W_i_* of an individual determines its probability of being selected to die in every time period. Note that this is different than Smolla and Akçay (2019) where selection was on fertility; i.e., the payoff due to culturally acquired traits affected the probability of getting selected to reproduce. Implementing fertility or mortality selection has no effect on our main results in homogeneous environments (see Fig. S15); all results discussed in the main text for mortality selection also carry over to fertility selection. However, in heterogeneous environments fertility selection can result in densely connected populations ‘getting stuck’ on a few traits, an ‘echo-chamber’ phenomenon as observed and discussed by Smolla and Akçay (2019). This effect makes densely connected populations unable to track changing environments.

The payoff *W_i_* in equation (3) is in principle unbounded, but in practice will be limited by the finite individual and social learning success of individuals. However, in heterogeneous environments with high trait variance, a few very valuable traits (very high *u_j_*) might end up having a disproportionate effect on survival in our simulations and drive artifactual results. To guard against such a nuisance outcome, we assume that fitness exhibits diminishing returns with utility. Specifically, we assume the probability of death is proportional to 1/*M*(*W_i_*), where *M*(*W_i_*) is a Michaelis-Menten function *M*(*W_i_*) = 1 + (*V*_max_*W_i_*)(*K* + *W_i_*)^-1^, with upper limit *V*_max_ = 50 and half rate constant *K* = 50, and a minimum payoff of 1. The high value of the half-rate constant means that for most of the parameter range, this probability increases almost linearly with Wi while guarding against unrealistic strong selection effects with extreme heterogeneity in trait payoffs.

A simulation turn ends with re-calculating each individual’s payoff. A new simulation round starts with the removal of an individual, and selection of a random survivor.

### 2.4 Population size

The effect of population size on cultural evolution has been somewhat controversial recently (see e.g. Fay et al., 2019; Henrich, 2004b; Powell et al., 2009; Shennan, 2001). We consider how population size affects the coevolution of culture and network structure by simulating populations of different sizes, *N* ∈ {25, 50, 75,100, 200}.

### 2.5 Simulation parameters

If not stated otherwise, we run all simulations with *N* = 100 and *M* = 500, for 5,000 generations (one generation is defined as *N* death-birth events, with data being averaged over the last 200 generations) and 200 repetitions, with mutation rate *m* = 0.05 (with SD = 0.05 for *p*_n_ and SD = 0.005 for *p*_r_), innovation success rate *α* = 0.02 and social learning success rate *β* = 0.5. Complex networks are initialised with random values drawn from uniform distributions (*p*_n_: *U*(0,1), *p*_r_: *U*(0, 0.1)). Individuals are initialised with empty repertoires.

### 2.6 Outcome variables

To compare differences in cultural knowledge between populations we record average repertoiresize 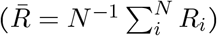, and mean highest per individual trait proficiency 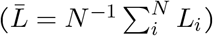. To compare evolved networks, we record degree centrality, and average weighted component size. Degree centrality, a measure for connectedness, is the average number of connections an individual shares with other individuals (higher degree centrality signifies more connections between individuals). As the emerging networks can have unconnected components, we calculate the average weighted component size as a measure for the component size distribution. We define average weighted component size as the sum of the product of the component size, *c_s_*, and the relative component size, 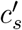 over all component sizes, that is 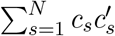. We calculate the relative component size 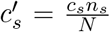, where *n_s_* is the number of components of size *s*, which is equivalent to 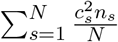. If not stated otherwise, all reported results are averages of the last 20% of generations.

## 3 Results

### 3.1 Homogeneous environments

#### 3.1.1 Two distinct pathways to cultural adaptations

We find that two distinct types of populations emerge from cultural selection in homogeneous environments (Fig. 1). The first type has low average connection probabilities (*p*_n_, *p*_r_) and features sparsely connected (low degree) networks with disjointed components (average component size less than *N*). The second type of population has high average connection probabilities and densely connected networks composed of a single connected component (average component size equals *N*; representative examples as insets in Fig. 1b). These populations display two distinct kinds of cumulative culture as characterised by the average repertoire size and average skill proficiency. Sparser networks only reach baseline proficiency on average, but have a slightly larger repertoire (Fig. 1c). In contrast, denser networks achieve higher average proficiency with only a slightly reduced average repertoire. Interestingly, we find that proficiency is high for a wide range of degrees, so long as the network is connected. In contrast, repertoire sizes are large for a wide range component sizes, as long as average degree is low and there are at least a few unconnected components.

**Figure 1.**
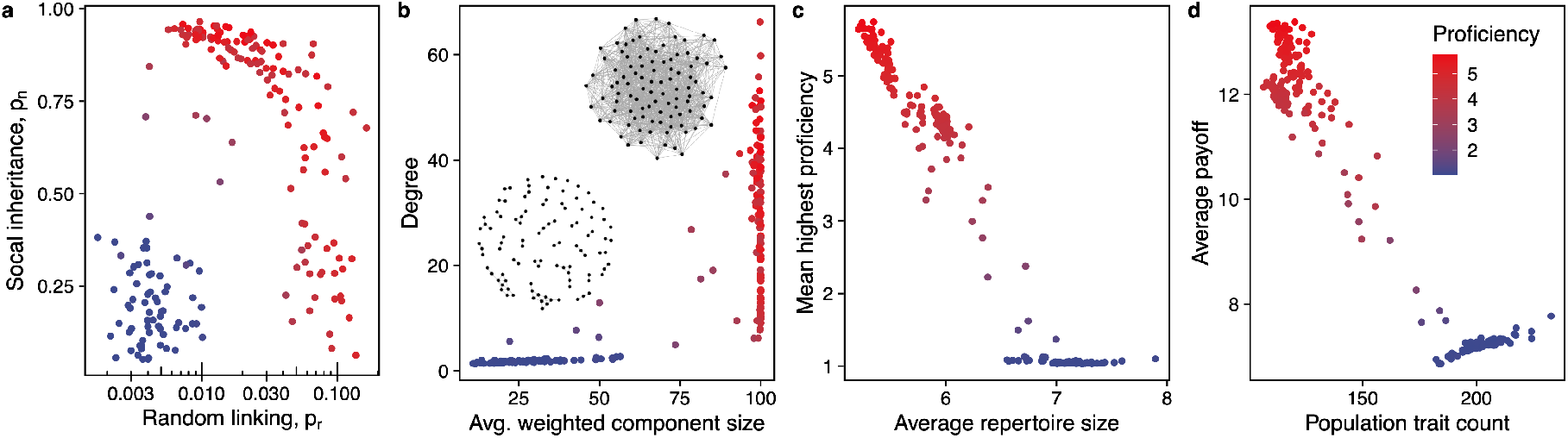
The two pathways to cultural adaptation. The four panels depict 200 populations from our simulations, with each panel showing different characteristics of the same set of populations. Each population is represented by a dot, coloured according to the mean proficiency of that population. When linking parameters evolve two distinct kinds of populations emerge: low vs. high mean linking traits (*p*_n_ and *p*_r_, panel a), low degree and small, unconnected components vs. high degree and a single connected component (panel b), large repertoires and low proficiency vs. smaller repertoires and high proficiency (panel c), and high trait diversity and low payoff vs. low trait diversity and high payoff (panel d). Simulations with *α* = 0.01, *β* = 1, *N* = 100, *M* = 500, running for 5,000 generations, *τ* = 0 and *σ* = 0.

While average repertoire sizes at the individual level differ only modestly between the two kinds of populations, the cultural makeup at the population level differs markedly. Populations with sparser networks possess almost twice as many unique traits compared to those with denser networks but they also have much lower average payoff at the same time (Fig. 1d). Further, the trait frequency spectrum differs. Populations with sparse networks have a much more even distribution with many traits at intermediate frequencies, whereas dense networks converge to a few traits known to almost every individual and a high number of traits at low frequencies (Fig. S2). The convergence to a few traits by the whole population allows these networks to increase proficiency in these traits.

#### 3.1.2 Low-payoff state persists due to cycling

Figure 1d shows that groups with larger repertoires have on average much lower payoffs than those with high skill proficiency. Why do high-repertoire populations persist in the long term despite this payoff disadvantage? Analysing simulation trajectories, we find that populations exhibit cycles in the *p*_n_-*p*_r_ space (Fig. S7) that repeatedly push them between the high-proficiency (and high payoff) and large-repertoire (and low payoff) states (videos of cycling behaviour available in SI). This cycling is a result of how different combinations of *p*_n_ and *p*_r_ relate to average payoff. Figure 2 shows the average linking parameters and associated average payoffs for 100 simulations over their last 100 generations. The density of the points in Figure 2 indicates where the populations spend their time in steady state. This plot highlights two important features: first, the average payoff stays high as long as either *p*_n_ or *p*_r_ or both are high. Second, there is a region separating the low- and high-linking probabilities that populations do not linger in. Plotting average payoffs across constant *p*_n_ transects in Figure 2b shows that this region corresponds to a ‘valley’ of low average payoff. Specifically, average payoff decreases with decreasing *p*_r_, but increases somewhat again at very low *p*_r_. Yet, time trajectories from our simulations show that populations consistently drop into this payoff valley from the high-payoff state (i.e., from the right-hand side with low *p*_n_) and emerge on the other side, in the low-payoff state (on the left-hand side, see also Fig. S8).

**Figure 2.**
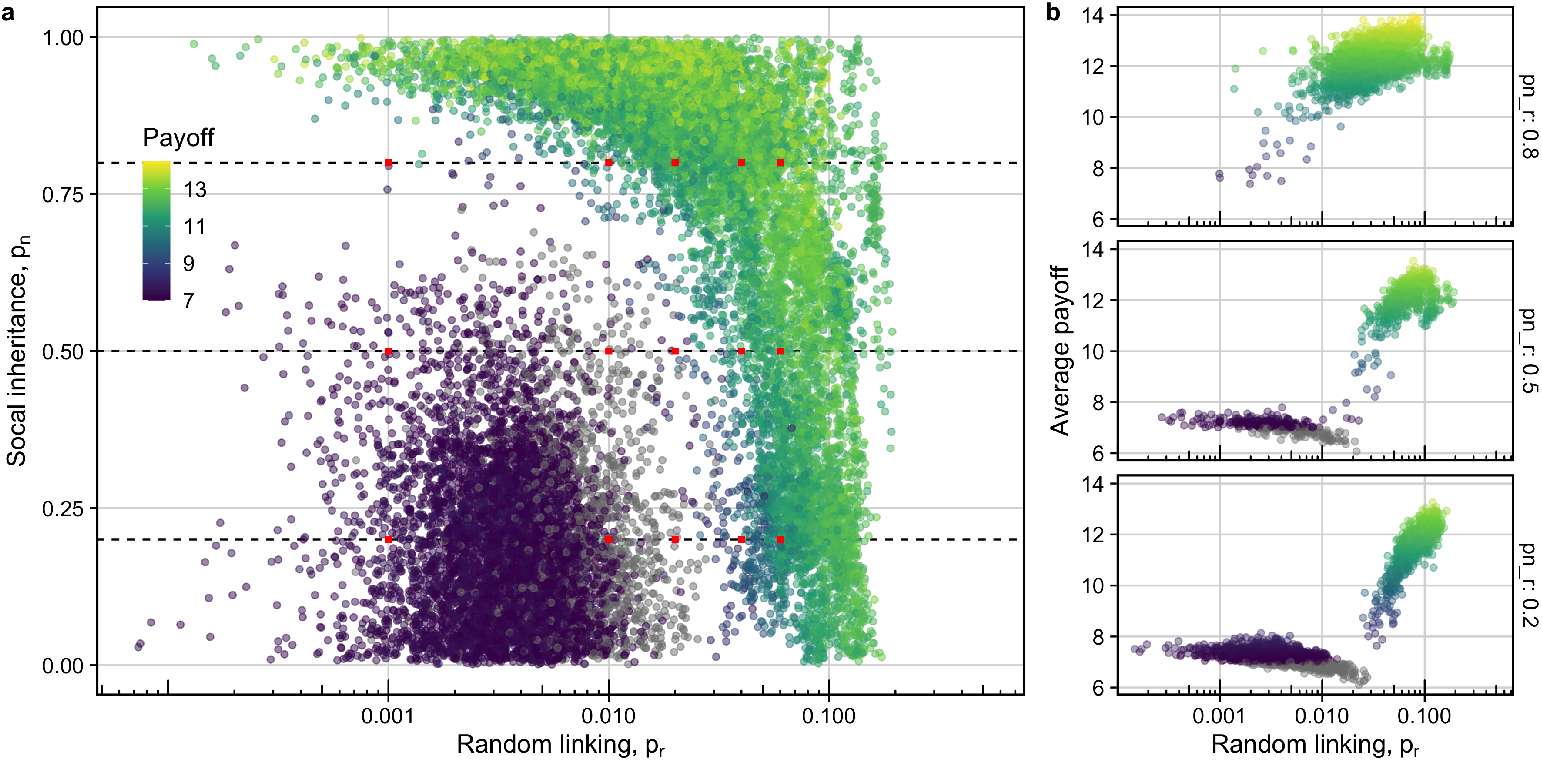
The fitness landscape for connection traits, *p*_n_ and *p*_r_. Panel a shows mean linking parameters and payoff for populations (one every 10 generations over the last 100 generations). The distribution of lighter and darker coloured points highlights the distinct population states separated by a fitness valley where the populations spend negligible time (for illustrative purpose payoffs < 7 shown in grey). Panels b-d depict cross-sections of this fitness valley at particular values of *p*_n_ (at the horizontal dashed lines in panel a). The red dots depict the resident *p*_n_ and *p*_r_ values used for Figure 3. Simulations with *α* = 0.01, *β* = 1, *N* = 100, *M* = 500, running for 5,000 generations, *τ* = 0 and *σ* = 0.

#### 3.1.3 Cycling results from the conflict between individual and group payoff

To explain this puzzling observation, we computed the local selection pressures acting on the linking traits *p*_n_ and *p*_r_ in populations that are fixed for a particular value of the linking traits and have the associated steady state cumulative culture. Figure 3 depicts the relative payoff (dis-)advantage of a mutation that changes the linking traits for an array of resident *p*_n_ and *p*_r_ values. This gives the local selective landscape at the individual level across the *p*_n_-*p*_r_ space and reveals a striking contrast between individual selection and population average payoff. Even though average payoffs at the population level are highest when either *p*_n_ or *p*_r_ or both are high, selection almost always favours reduced linking at the individual level.

**Figure 3.**
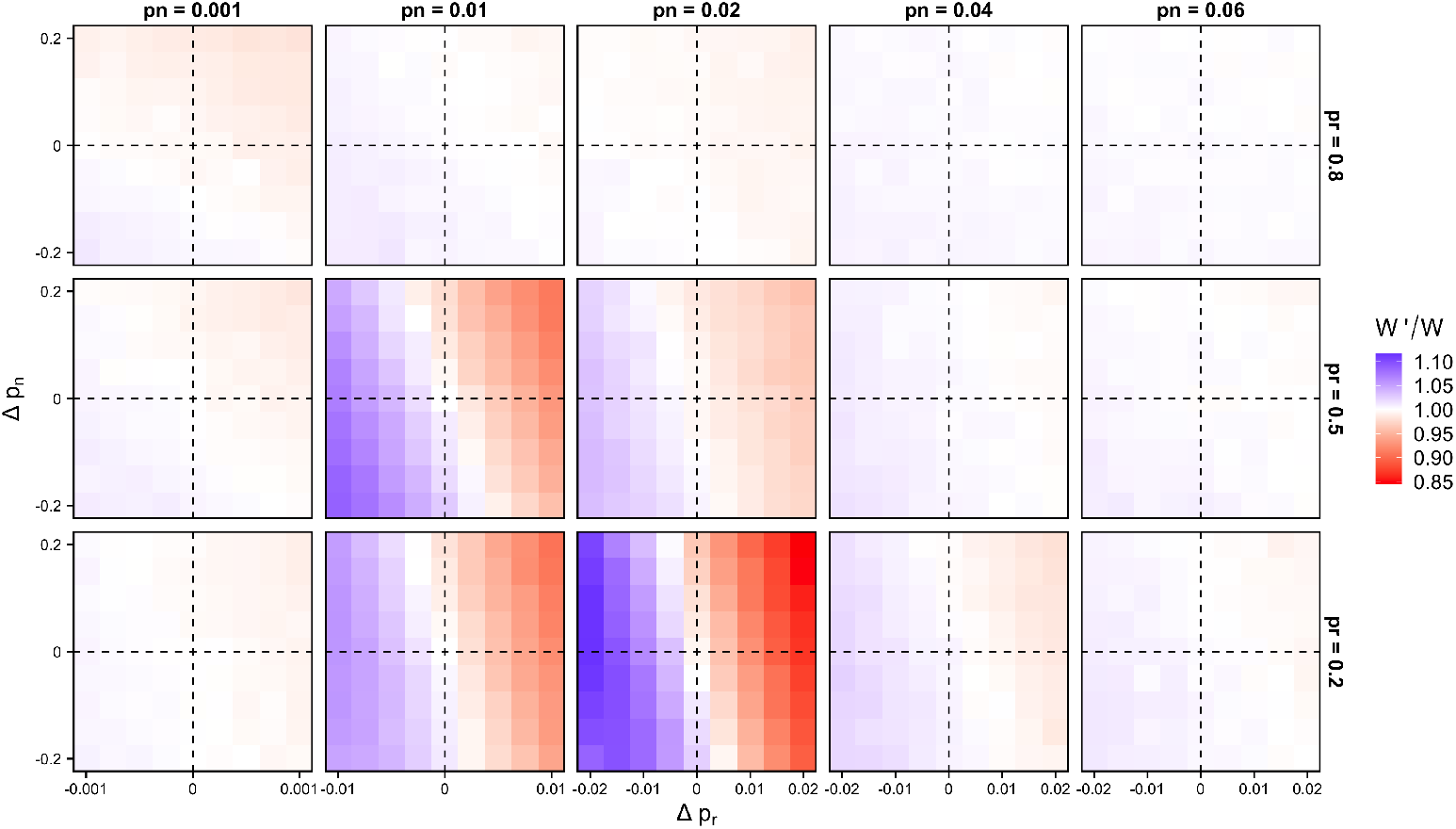
Local selection pressures on the linking traits. Each panel corresponds to results from ten populations fixed for a particular value of *p*_n_ and *p*_r_ (given on the right-hand side and top of the grid, respectively; these correspond to the red dots in Figure 2). For each panel we initialise 10 populations by running our cultural selection model with fixed connection traits and allowing the cumulative culture to come to a steady state. Then we introduce a single mutant that deviates from the resident linking parameters by Δ*p*_n_ and Δ*p*_r_, which are depicted on the x- and y-axes of each panel, respectively. Next, we calculate the relative payoff *W*’/*W* of the mutant relative to the mean payoff of the residents, as a result of the cultural traits the mutant learns and innovates. We repeat this 500 times for every combination of Δ*p*_n_ and Δ*p*_r_. Relative fitness is generally higher with lower *p*_n_ and *p*_r_, revealing individual level selection for disconnecting, despite the population level consequences for group level payoff depicted in Figure 2. Simulations with *α* = 0.01, *β* = 1, *N* = 100, *M* = 500, running for 5,000 generations, *τ* = 0 and *σ* = 0.

This discord between average group payoff and individual-level selection emerges from the interaction between emergent cumulative culture and social learning as we model it. Recall that high-payoff populations have dense networks and strong cultural convergence (that is, a small number of traits that are present in all repertoires and for which there is high proficiency). At the same time, because every individual also innovates random traits with a small probability, being connected to many others not only brings the individual in contact with the most common traits but also with those that are rare. However, because learning in our model requires repeated observations of a trait, the presence of a large number of rare traits reduces the overall success of social learning. This ‘distraction’ effect of rare traits applies most strongly to other rare traits (as common traits are observed at the same overall frequency no matter how many connections are made). Therefore, individuals with lower p_n_ and/or *p*_r_ (who on average connect to fewer others) will be more likely to learn more socially and have a higher payoff. This individual benefit drags the population towards the valley in the average fitness, and ultimately to the low-payoff state. Figure 3 shows that the strongest pull towards the low payoff population state occurs in the fitness valley described above. Populations can escape from this sparsely connected, low-payoff state either by drift, due to the weak selection with low *p*_r_, or by trading random connections for socially inherited ones: at low *p*n and *p*_r_ (lower left panels in Figure 3), individual selection can favour increased *p*_n_ as long as it is combined with decreased *p*_r_. This is because an increase in social inheritance links the individual to a clustered neighbourhood that is more likely similar in their trait repertoire. This creates local cultural convergence and reduces the number of unique rare traits an individual is exposed to. These clustered components can thus achieve higher payoffs and grow to take over the population, a kind of component-level selection that takes populations back to the connected, high-proficiency state.

Transitions between high and low payoff state happens more often in smaller populations than large populations (see Table S1) and larger populations spend more time in the high payoff state in the long run. However, even relatively large populations (*N* = 500) can spend 30% of time in the low-payoff state given our base parameters.

This cycling behaviour is a previously unrecognised example of a social dilemma where the interest of the individual is at odds with that of the group. The group benefits from being highly connected, as this allows for cultural convergence and subsequently an increased skill proficiency. While the individual benefits from this situation, they benefit even more by reducing their linking to the group. However, ultimately, this results in the social networks breaking apart and a loss of accumulated proficiency that disconnected networks cannot produce or sustain. This is a new kind of social dilemma between cultural adaptation at the group level and selection for the social structure that supports it at the individual level. Crucially, the assumption that social learning requires repeated exposures is necessary for these results, since this assumption creates a trade-off between overall success of social learning and the number of traits. If social learning can follow from single exposures, there is no such trade-off, and accordingly, populations evolve to be highly connected but without the convergence on a few traits and associated high proficiency (see Figures S9 and S17).

#### 3.1.4 Frequent innovation selects for sparser networks and lower payoff

Next, we explore how individual innovation and social learning rates influence the coevolution of network structure and cumulative culture. If there is no social learning (*β* = 0), increasing innovation rate α always increases average payoff (Fig. 4a), independent of the environment. This is straightforward because without social learning, an individual’s repertoire and proficiency depends solely on innovation, and more innovation means both larger repertoire size and higher proficiency. Social network structure, as expected, evolves neutrally in this case (Fig 4b).

**Figure 4.**
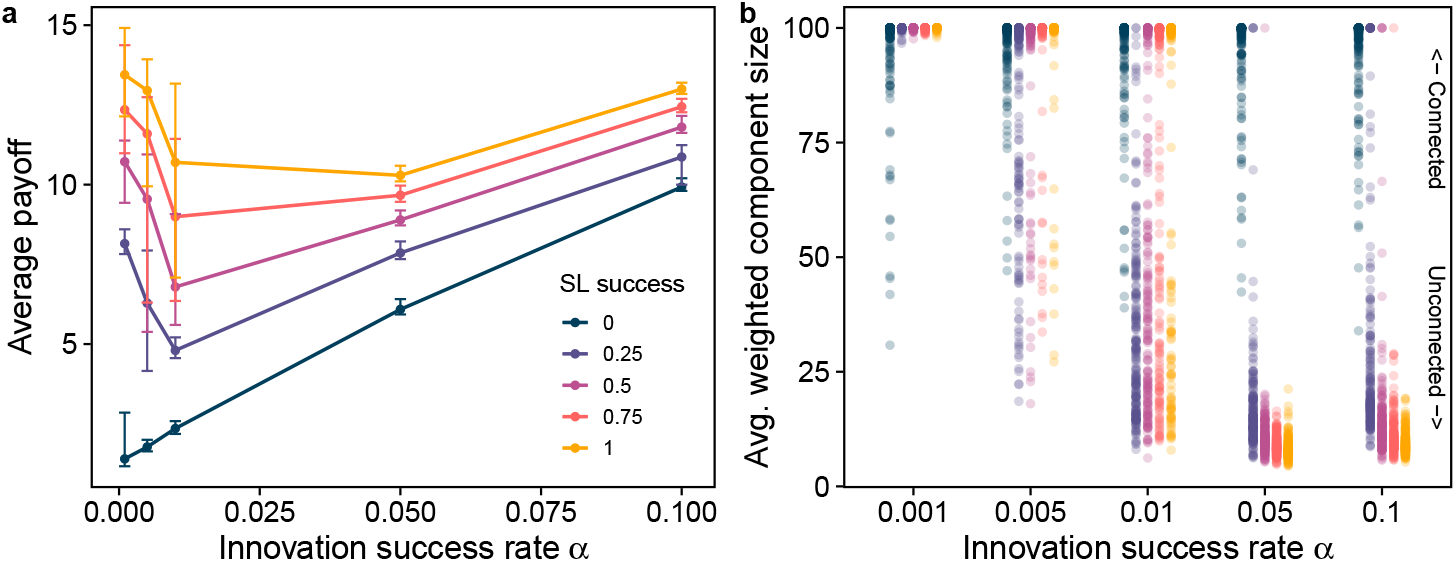
The rate of innovation and social learning affects payoffs (a) and social network structure (b). Panel a depicts average payoffs as a function of innovation success rate *α* for different social learning success rates (*β*). In the absence of social learning (*β* = 0), we find that increasing innovation always increases payoffs. However, with social learning (*β* > 0), we find that average payoffs decrease with innovation rate initially before recovering at high innovation rates. Panel b depicts the average weighted component size, a measure of the connectivity of the network, with social learning success rate *β*, for different values of individual learning rates. It shows that as long as there is any social learning (*β* > 0) higher individual innovation rate results is less connected networks. Simulations with *N* = 100, *M* = 500, running for 5,000 generations. Error bars in a represent 90% confidence intervals.

In the presence of social learning, however, we again find an unexpected result: increasing individual innovation rate *α* decreases the average payoffs of the population (Fig. 4a). This counter-intuitive pattern again happens because of the interplay between network structure and social learning: when innovation rates are low, individuals carry few or no rare idiosyncratic traits. Therefore, connecting to more individuals does not carry a cost in terms of reduced learning probabilities, which reduces the individual level selection to disconnect. Populations thus remain highly connected, converge on a few traits, and retain improvements in proficiency in these even if they are very rare. As individual innovation rates increase however, each individual innovates more traits, which in a highly connected population reduces the probability to acquire traits through social learning. Therefore, the individual level selection for reduced connections becomes stronger, and as a result, populations are more likely to spend time in the sparsely connected, low payoff state (Fig 4b). As innovation rates increase further, populations spend all their time in the sparsely connected state, and average payoff only recovers when innovation rates are so high that individual innovation starts compensating for the lack of socially learned traits. Note that increased individual innovation rate is always directly beneficial for the innovating individual (as it simply increases its proficiency or repertoire) but our results illustrate it can inhibit accumulation of culture because of its effect on network evolution, another instance of a conflict between individual selection and group-level cultural adaptation.

### 3.2 Heterogeneous environments

#### 3.2.1 Environmental turnover and trait variance affects the networks that evolve

In heterogeneous environments, where payoffs vary across traits and time, we find that the variance in trait payoffs and the turnover rate of payoffs play a crucial role in determining the network structure and associated cumulative culture. Specifically, high proficiency culture evolves much more readily in stable environments with highly skewed utility distributions (Fig. 5a), which coincides with occurrence of connected graphs (Fig. 5b). In contrast, environments with fast environmental turnover but highly variable utilities favour the evolution of sparse networks with larger individual skill repertoires (Fig. 5b) and a larger cultural repertoire (Fig. 5d). Here, coordination on a few traits and accumulation of proficiency does not happen fast enough before payoffs change again. These differences disappear as utility distributions become increasingly uniform (small *σ*). That said, similar to the homogeneous environment case, we find both kinds of networks and their associated repertoire type (Fig. 5e) in the majority of environments that we tested. Only where utilities are highly heterogeneous and environments are stable, the low payoff state is absent.

**Figure 5.**
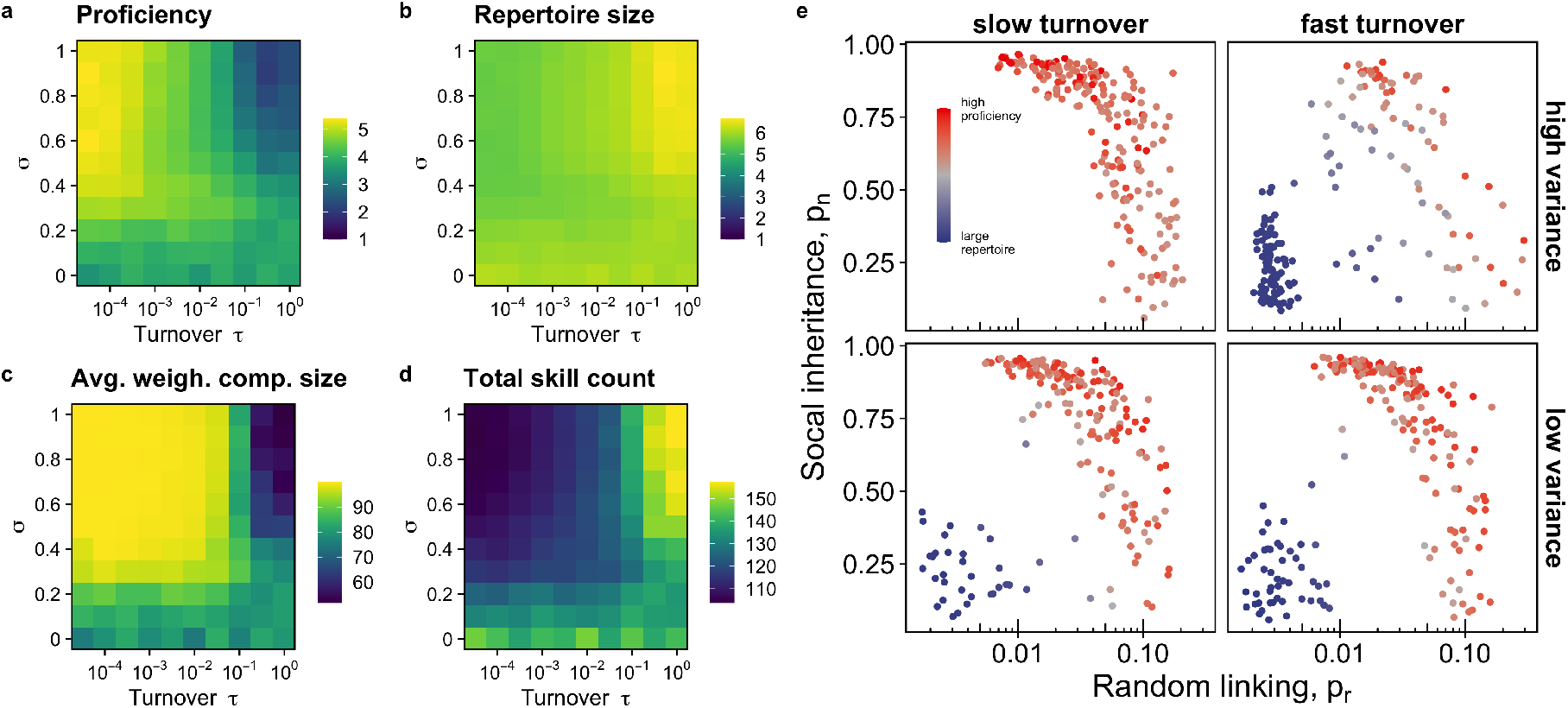
Environmental heterogeneity affects the viability of the generalist and specialist repertoire populations. When linking parameters evolve in different environments, we find stronger reliance on high proficiency (and there are fewer traits in the population overall) where turnover is low and utilities are highly skewed (large *σ*), whereas in environments with frequent turnover we find stronger reliance on larger repertoires. Simulations with *α* = 0.01 and *β* = 1, *N* = 100, *M* = 500, running for 5,000 generations, averaged over 200 repetitions. False colour scale in e is based on the following data transformation of average repertoire size (R) and highest proficiency (L): *L*/max(*L*) – *R*/max(*R*). Subset of data in e with *τ* ∈ {10^-3^, 1} and *σ* = {0.2,1}. See Fig. S10 for additional results.

#### 3.2.2 Larger populations accumulate more proficiency in stable environments

Finally, we vary population size and find that as populations become larger they are more likely to accumulate higher proficiency and be more connected (Fig. S12). However, this feature is conditional on environmental stability. Where utilities change frequently, we do not find an effect of population size: larger populations stay disconnected and in the low proficiency, broad repertoire state. It is interesting to note that the cumulative nature of learning in our model allows individuals to reach higher proficiency in larger populations. However, this is not the case for repertoire size. In fact, individuals in unconnected graphs of large populations achieve similar if not smaller repertoire sizes compared to smaller populations (Fig. S14).

## 4 Discussion

Our results uncover some fundamental tensions in the coevolution of cumulative culture and social network structure through which cultural traits are propagated. We demonstrate two distinct pathways to cultural adaptation is available to populations in a setting where both learning new traits and increasing proficiency in existing traits is available. These two pathways (broad repertoire versus high proficiency) correspond to distinct population structures and cumulative culture features. At the same time, we uncover two general conflicts populations face when navigating these pathways. First, we show that although groups do best on the aggregate when they are well-connected, coordinate on a few core cultural traits, and build up proficiency, selection favours disconnection at the individual level. This creates a previously unrecognised type of conflict between individual and group-level payoff. Second, we show that increasing individual innovation rates results in less connected networks and a lower population level payoff. This is because all else being equal, higher individual innovation increases the trait diversity around an individual which again makes social learning less successful, and increases the selection against social connections. Since individual innovation is also individually beneficial (as it only results in individuals knowing more traits or having higher proficiency) this is another dimension of conflict between individual and group-level payoff. It is important to note that both of these results obtain despite there being no inherent cost to making or maintaining social connections.

Both of these conflicts between group- and individual-level benefits arise from the nature of social learning. Our model does not inherently privilege learning or innovating new traits over increased proficiency in an existing trait. However, individuals can only learn socially when they are repeatedly exposed to the same trait. Having more connections exposes an individual to more idiosyncratic traits that have been individually innovated, which reduces the probability of successful social learning overall. Having a higher innovation rate similarly reduces the social learning success of one’s social connections. The reduction in overall social learning probability with the number of options available is akin to the well-documented phenomenon of “choice overload” in consumer psychology (Chernev et al., 2015; Iyengar and Lepper, 2000; Scheibehenne et al., 2010): having too many choices for a consumer good can result in a number of counterintuitive effects, including a reduction in the probability of purchasing anything at all. In our case, this pattern flows directly from the requirement that in order to socially acquire a trait, individuals have to be exposed to it multiple times. This means that the probability of social learning is non-linear in the frequency of the trait in an individuals social neighbourhood and decreasing in the number of skills that are available for observation.

Because of this conflict between group-level payoffs and individual level selection, high proficiency culture and the social structure that maintains it can be considered a public good: maintaining the social learning infrastructure for high proficiency culture requires individuals to make more connections and innovate less than what would maximise their own payoffs. As in other public goods, groups that can counteract socially harmful individual incentives. For example, Ache hunter-gatherers maintained a high yearly interaction rate and high number of total connections for their population size compared to the Hadza hunter-gatherers in part by organising inter-band club fight rituals (Hill et al., 2014). Likewise, in Southern India, participating in religious rituals results in denser connections (Power, 2018). Our results provide a novel hypothesis for the evolution of rituals and social norms that promote social connections can enforce connectivity, cultural convergence, and resulting high proficiency, which can have an advantage in competition with other groups. One further factor modulating the connectivity of the learning network might be the traits themselves: traits used in cooperative foraging or social rituals will necessarily have more connected learning networks than traits that are mostly for personal or household use, as knowledge about medicinal plants (Salali et al., 2016).

Likewise, as we find too much individual innovation ultimately is detrimental to high proficiency culture, we might expect norms that enforce social learning at the expense of individual innovation, even if there is no inherent difference between the traits that are socially transmitted. Such normative behaviour discouraging individual innovation has been described, for example, for the transmission of pottery skills among three ethnic groups (Dii, Duupa, and Doayo) in Cameroon (Wallaert-Pêtre, 2001). Here, practitioners form tight communities that limit access to knowledge and reject departures from socially admitted norms. This leads to highly conserved methods and products, which is often observed in the context of formal apprenticeships (see e.g. Lancy, 2012). Similarly, Buckley and Boudot (2017) provide an example of loom and weaving technology in Southeast Asia, where knowledge is primarily transmitted form mothers to daughters over a long apprenticeship, which similarly leads to relatively low rates of innovation.

Another potential solution to the problem of maintaining a specialised and high proficiency cumulative culture is role-specialisation. Our model considers a simple, undifferentiated population; this might be an ancestral state for human populations but the tensions identified in our model are expected to impose strong selection for differentiation of roles with respect to social learning. Specifically, high effective connectivity with respect to social transmission of traits can also be achieved if learning happens mostly through a subset of individuals that specialise in teaching social information. These individuals could also invest into making transmission more efficient by directed demonstrations or focusing attention (Ventura and Akcay, 2022). This is an additional selective advantage for the evolution of teaching, in addition to ensuring transmission fidelity (Castro and a Toro, 2014).

Our results with heterogeneous traits and environmental turnover highlight an additional dimension of complexity in the coevolution of cumulative culture and social structure. Environmental turnover rate emerges as an important modulator of the type of network structure and cumulative culture, with rapid environmental turnover favouring disconnected graphs, larger repertoires, and low proficiency. These results are in line with previous results (Deffner and Kandler, 2019; Kolodny et al., 2015b) showing that stable environments lead to more specialised kinds of culture. Consistent with our predictions, Kalan et al. (2020) found that chimpanzee groups living in more variable environments exhibited greater trait diversity. On the other hand, heterogeneity in payoffs at a given point in time has contrasting effects depending on the turnover rate: at low turnover rates, where payoffs update infrequently, larger variance favours populations that find the high payoff traits and increase proficiency in them by becoming well-connected. However, at high turn over rates it becomes likely that a high payoff trait will no longer be high payoff by the time proficiency is accumulated, which forecloses this route to cultural adaptation. Therefore, individuals are selected to increase their repertoire size to try to maximise the probability of learning at least some high-payoff traits. Notably, in most cases, populations still visit both the highly connected, high proficiency state and the sparsely connected, broad repertoire state, indicating that environmental heterogeneity and turnover modulate but not entirely eliminate the tensions described above.

Our model highlights how the interaction of individual innovation, social learning, and connectivity creates emergent constraints on the evolution of cumulative culture. This has implications for the much-discussed topic of how cultural complexity relates to population size. We find that cultural complexity does not keep increasing with population size (Figure S12 because further increases in complexity run into the limits of social learning at the individual level. This effect was also observed in a previous model by Mesoudi (2011) where social learning and individual innovation both were assumed to incur a direct cost, putting a limit to the amount individuals can learn. These results suggest that a population size effect on cumulative culture will be limited in the absence of further innovation in learning technology (as discussed above) that can ameliorate the trade-offs in social learning. We further show here that the effect of population size on cumulative culture is contingent on the environment: the increase in cultural complexity in the form of increased proficiency requires that the environment change relatively slowly. Otherwise, what is profitable changes before any proficiency can be accumulated through cultural selection, and population size has no effect on the cultural complexity.

Overall, our model points to previously unappreciated tensions between different pathways to cultural adaptation in groups and the incentives at the group- and individual-level that arise from the coevolution of interaction networks and cumulative culture. Resolving these tensions in different ecological settings likely played an important role in human social evolution and our ability to do so might be one reason for the unique success of our species.

## Acknowledgement

The authors thank Bryce Morsky, Andrew Tilman, Laurel Fogarty, and Colin Twomey for their comments on the project.

## Code availability

All simulation code used in this paper is available at https://github.com/marcosmolla/pathways_cultural_adaptation.

## Author contributions

E.A. secured funding. M.S. and E.A. contributed to the study design, M.S. conducted analyses, M.S. wrote the initial draft of the manuscript and all authors contributed to revisions.

## Additional information

Supplementary material is available for this paper at https://github.com/marcosmolla/pathways_cultural_adaptation.

## Supplementary Material

### S1 Payoff distribution

To simulate heterogeneous trait utilities, we draw random values from a lognormal distribution. Examples for relevant values in the main text are shown in Fig. S1.

**Figure S1.**
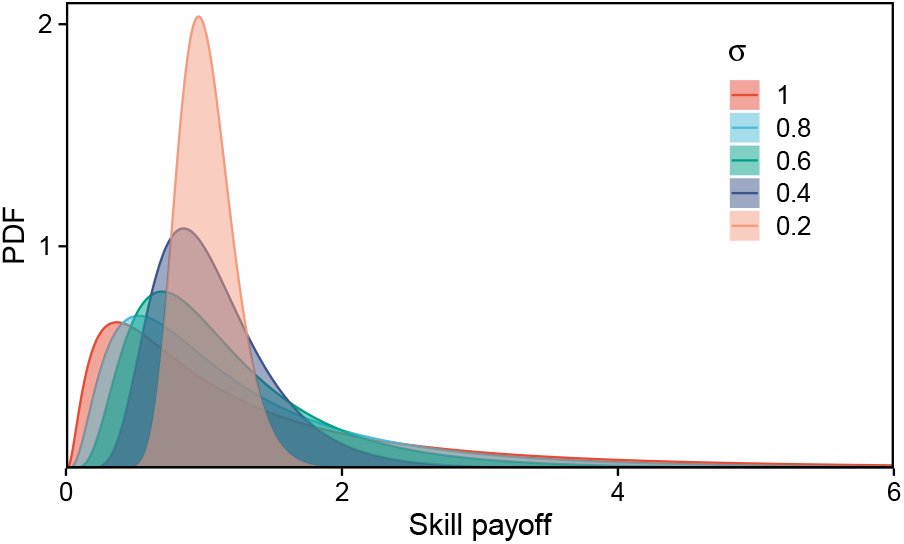
Probability density function for the different payoff variances used in the main text. Values are drawn from a lognormal distribution, Lognormal (*μ, σ*^2^) with *μ* = 0 and *σ*^2^ ∈ {0.2, 0.4, 0.6, 0.8,1}. As *σ*^2^ increases, the distributions becomes more skewed. Payoffs drawn from these distributions have a mean of *μ* = {1,1.1,1.2,1.4,1.6} and a variance of var = {0.04, 0.20, 0.62,1.7,4.66}.

### S2 Fixed homogeneous environments

#### S2.1 Cultural selection on complex graphs

Next, we let the linking parameters evolve in fixed and homogeneous environments. We observe which networks emerge, and how this affects individual and population-level culture.

**Figure S2.**
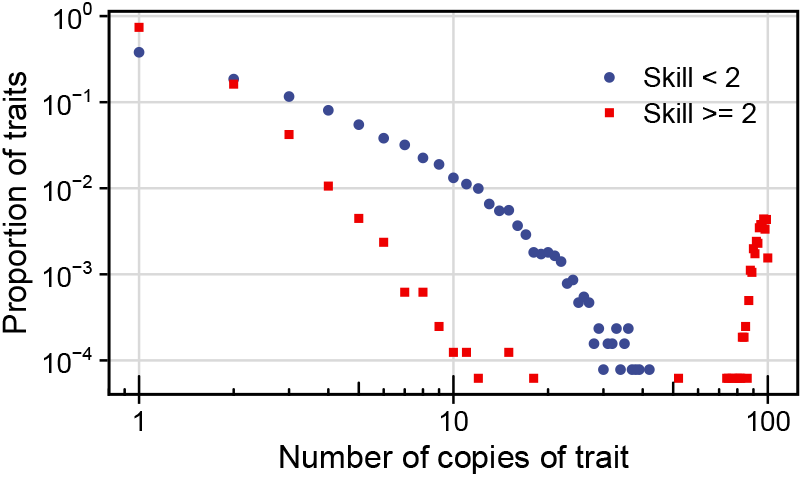
Trait frequency distributions differ between high proficiency and large repertoire populations. Populations with low average skill have in general more traits with intermediate frequency, whereas those with higher skill have a few traits with very high frequency in the population. Simulations with *α* = 0.01 and *β* = 1, *N* = 100, *M* = 500, running for 5,000 generations, *τ* = 0 and *σ* = 0.

#### S2.2 Robustness checks

We ran several robustness checks. First, to test that selection is acting here, we remove selection. When the two linking parameters are left to drift, we do not observe these two kinds of populations. Instead we find networks that are mostly connected, and populations with high proficiency but none with large repertoires (Fig. S3). Next, we ran simulations with fixed linking parameters to test that the differences in repertoire size and proficiency are, in fact, a result of the social network (Fig. S4). We observe patterns that correspond with the above results. Finally, to ensure that the observed patterns do not rely on the complex networks that we simulate here but are a general phenomenon, we simulated learning on random graphs with different degree. Consistent with the results above we find that repertoire size and count of unique traits decrease and proficiency and payoff increase as degree increases (Fig. S5,S6).

##### Neutral selection

For the neutral case (Fig. S3), we see that *p*_n_ and *p*_r_ spread equally in all directions. This results in networks that are mostly connected (average weighted component size equals *N*). We also see that there is a proficiency and repertoire size trade-off, as we would expect. Populations with connected networks also have on average high skill proficiency, albeit smaller repertoires and fewer overall known traits.

**Figure S3.**
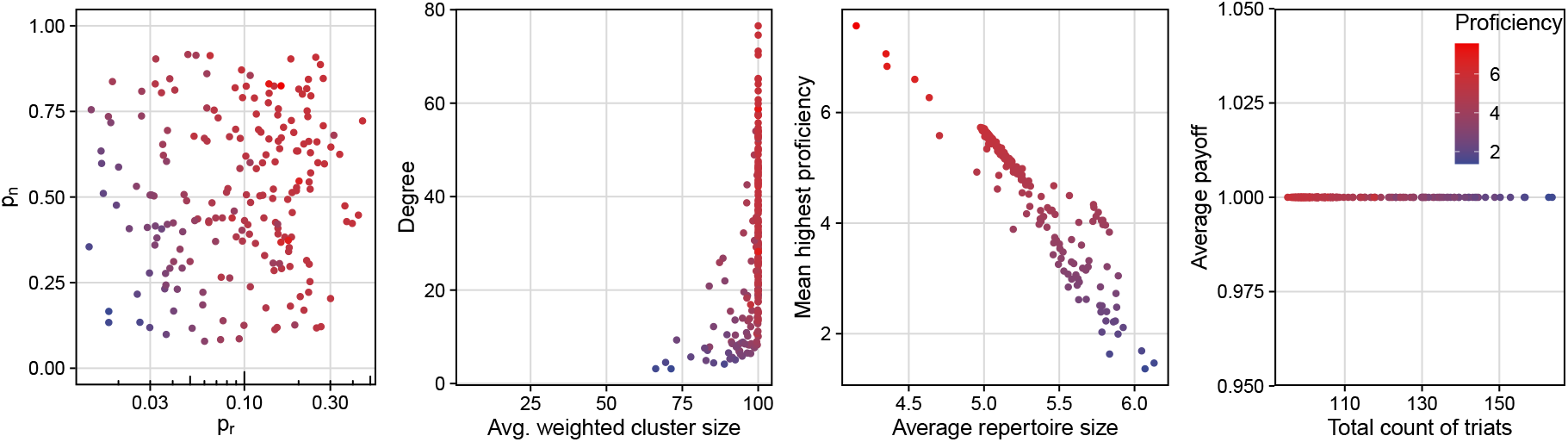
The neutral evolution of linking parameters *p*_n_ and *p*_r_ results in mostly connected graphs (large component size). In all plots colour indicates mean highest proficiency of a population. We find that populations with larger average degree reach higher proficiency. Populations with these networks also know on average fewer different traits. There is a clear negative relationship between proficiency and repertoire size, which is based on the trade-off during the limited number of learning turns: learn a new trait versus improve a known trait (see also Smolla and Akçay (2019)). Simulations with *α* = 0.02 and *β* = 1, *N* = 100, *M* = 500, running for 5,000 generations, *τ* = 0 and *σ* = 0, averaged over 200 repetitions.

##### Cultural selection on complex graphs with fixed linking parameters

We find two repertoire types with according network shapes when we fix linking parameters in simulations (so, there is still selection on survival, but linking parameters do not change over time). High *p*_n_ as well as high *p*_r_ lead to connected graphs, which result in a low overall skill count and small average repertoire sizes but higher skill level.

##### Cultural selection on random graphs

We run the learning algorithm described in the Methods section on random graphs (Fig. S5 and S6) to provide another reference point for the simulation with complex networks in the main text. We simulate different population sizes as well as different average number of neighbourhoods an individual is sampling from (so in principle these are random graphs).

**Figure S4.**
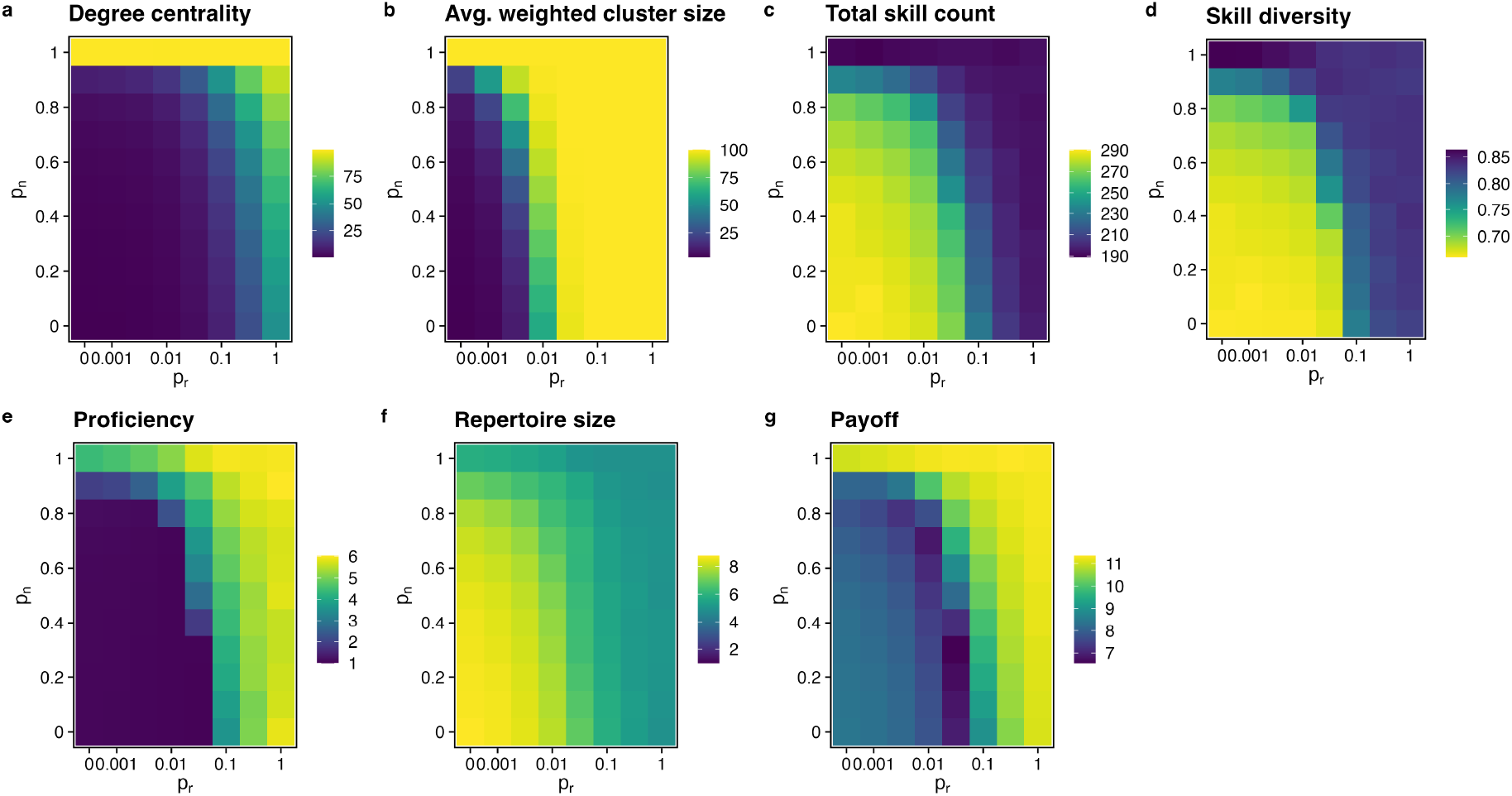
Linking parameters affect whether populations rely on high proficiency or large repertoires. The space that the two linking parameters *p*_n_ and *p*_r_ span can be separated into two areas. First, populations with low *P*_n_ and low *p*_r_ form networks with lower degree centrality (a) that are largely unconnected (b). They have the most diverse trait knowledge (c), which also means that they have the lowest proficiency (e) albeit the largest individual repertoires (f). The second zone is for high *p*_n_ and/or high *p*_r_, where we find connected graphs with high degree centrality. Here, proficiency is high and repertoires are small. Overall, populations in this zone know a smaller number of traits in total, however, they have on average a higher payoff then populations in the other zone. Simulations with *α* = 0.02 and *β* = 1, *N* = 100, *M* = 500, running for 5,000 generations, *τ* = 0 and *σ* = 0, averaged over 200 repetitions.

For the Proficiency/Repertoire size space, we find two types: low and high proficiency, whereby the latter has a slightly smaller repertoire size on average. Also, larger networks have on average higher proficiency and smaller repertoires when compared to smaller networks. Though, larger networks on average have a larger number of total traits.

**Figure S5.**
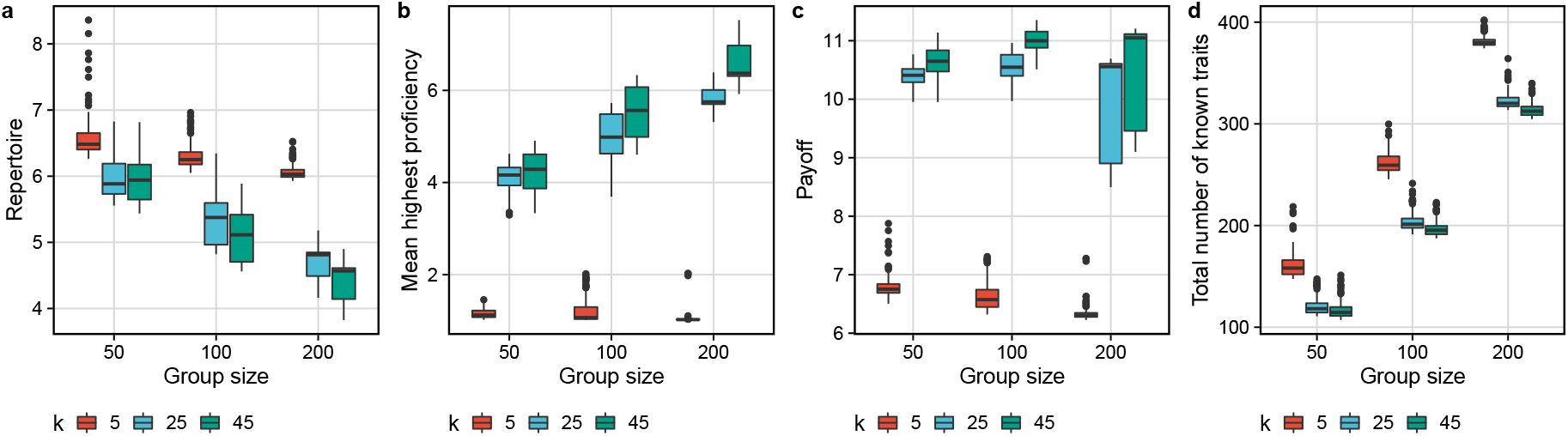
Population size as well as degree centrality affect individual and population-level culture. Individuals in larger populations have on average smaller repertoires and higher proficiency. Larger populations also know on average more different traits than smaller populations. Across all population sizes, individuals in networks with smaller *k* have larger repertoires and lower proficiency, and yield a lower payoff. Populations with smaller degree also know on average more different traits than those with a higher degree. Payoffs on average increase with population size, at least for high degree centrality. However, for low degree payoff decrease for larger populations. This is because of the larger number of known traits, which causes there to be a very high trait diversity in any individual’s neighbourhood which ultimately leads to many individuals learning even less (which we can see in the smaller average repertoire and the lower average proficiency of the large groups). We simulate populations with complete graphs but allow individuals only to learn from *k* random neighbours.Simulations with *α* = 0.02 and *β* = 1, *N* ∈ {50,100, 200}, *M* = 500, running for 5,000 generations, *τ* = 0 and *σ* = 0, with 200 repetitions for each set of parameters.

**Figure S6.**
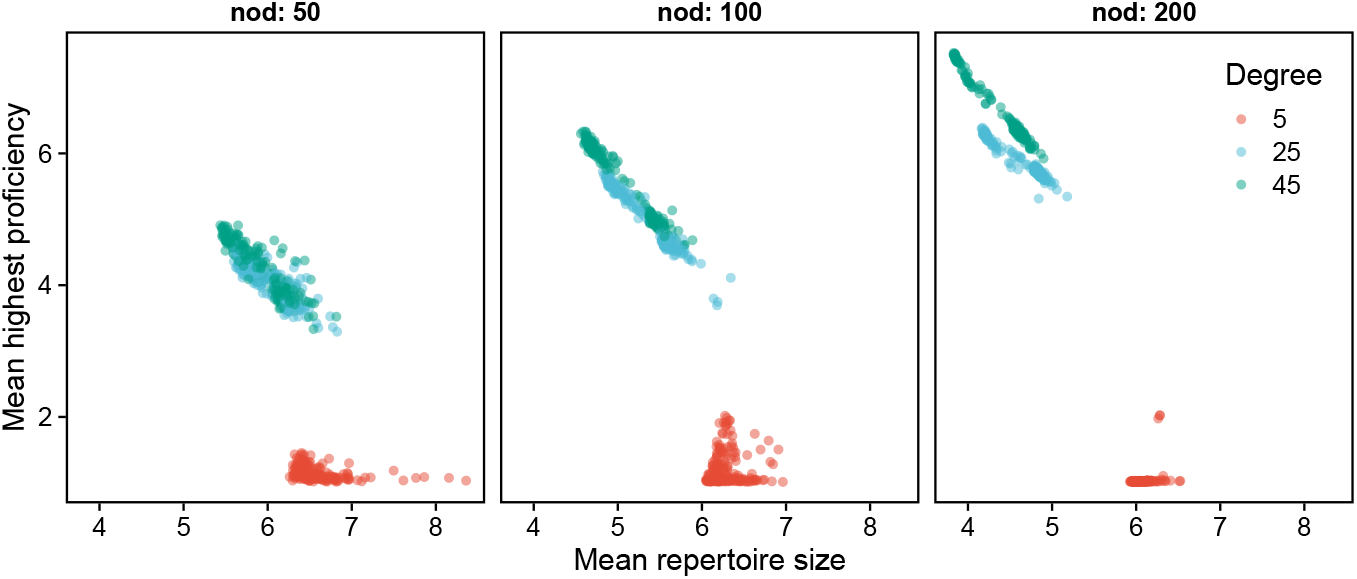
Connectivity affects repertoire type. Populations with higher average degree rely on high proficiency, whereas populations with sparser networks have low proficiency but slightly larger repertoires. As population size increases, so does the average highest proficiency reached in populations with intermediate and high degree centrality. Proficiency learning benefits from more frequent innovation in larger populations and cultural convergence due to high connectivity. In contrast, larger populations with sparse networks do not achieve larger repertoires. Despite harbouring more unique cultural traits (Fig. S5d), there is still a limit to the diversity in an individual’s neighbourhood for successful social learning. In fact, the larger number of cultural traits in the population leads to a slightly reduced repertoire size and consequentially smaller average payoff (Fig. S5c). Data from simulation in Fig. S5.

**Table S1.** Results of Markov chain analysis of cycling behaviour. Using data from simulations with different population sizes *N* (all with homogeneous environments, *τ* = 0, *σ* = 0, running for 40,000 generations, with *α* = 0.01 and *β* = 1), we first counted the number of times populations switched between the high and low payoff state. That is, we counted the number of times when a population’s mean payoff exceeded 60% of the maximum payoff from across all simulations, or fell below this threshold (we tested different thresholds but found this threshold to best separate the high and low payoff simulations). Next, we pooled the data from all 100 simulations and calculated the probability for a switch to occur in a single population from one generation to the next. We used these probabilities to construct a transition matrix for a Markov process and calculated the stationary distribution of both states (high and low payoff).

| | $N = 50$ | $N = 100$ | $N = 500$ |
| --- | --- | --- | --- |
| Prob. switch Up to Down | 0.01013 | 0.000114 | 2.10e-05 |
| Prob. switch Down to Up | 0.01018 | 0.000141 | 4.73e-05 |
| Stationary proportion high | 0.5 | 0.55 | 0.69 |
| Stationary proportion low | 0.5 | 0.45 | 0.31 |

#### S2.3 Cycling

Figure S7 shows trajectories of five example populations at different time points, illustrating the cycling phenomenon described in the main text. Figure S8 provides a closer look at the transitions between high and low payoff states.

#### S2.4 Simple contagion based social learning

Here, we show that the social learning model that requires repeated exposures is necessary to get the two distinct pathways. The repeated exposures requirement underlies our assumption that the probability of socially learning a trait is proportional to the square of its frequency in an individual’s neighbourhood. Figure S9 shows results of simulations in homogeneous, fixed environments where instead we assume social learning probability is proportional to the frequency of the trait in the neighbourhood. It shows that populations are selected to be always well connected, accumulate very high repertoires compared to the repeated exposure model and low proficiencies. The latter happens because in the absence of coordination on a few traits, more innovations that improve the proficiency get lost before they can get transmitted. This shows that the conflict between individual selection and group selection is absent with single exposure social learning.

**Figure S7.**
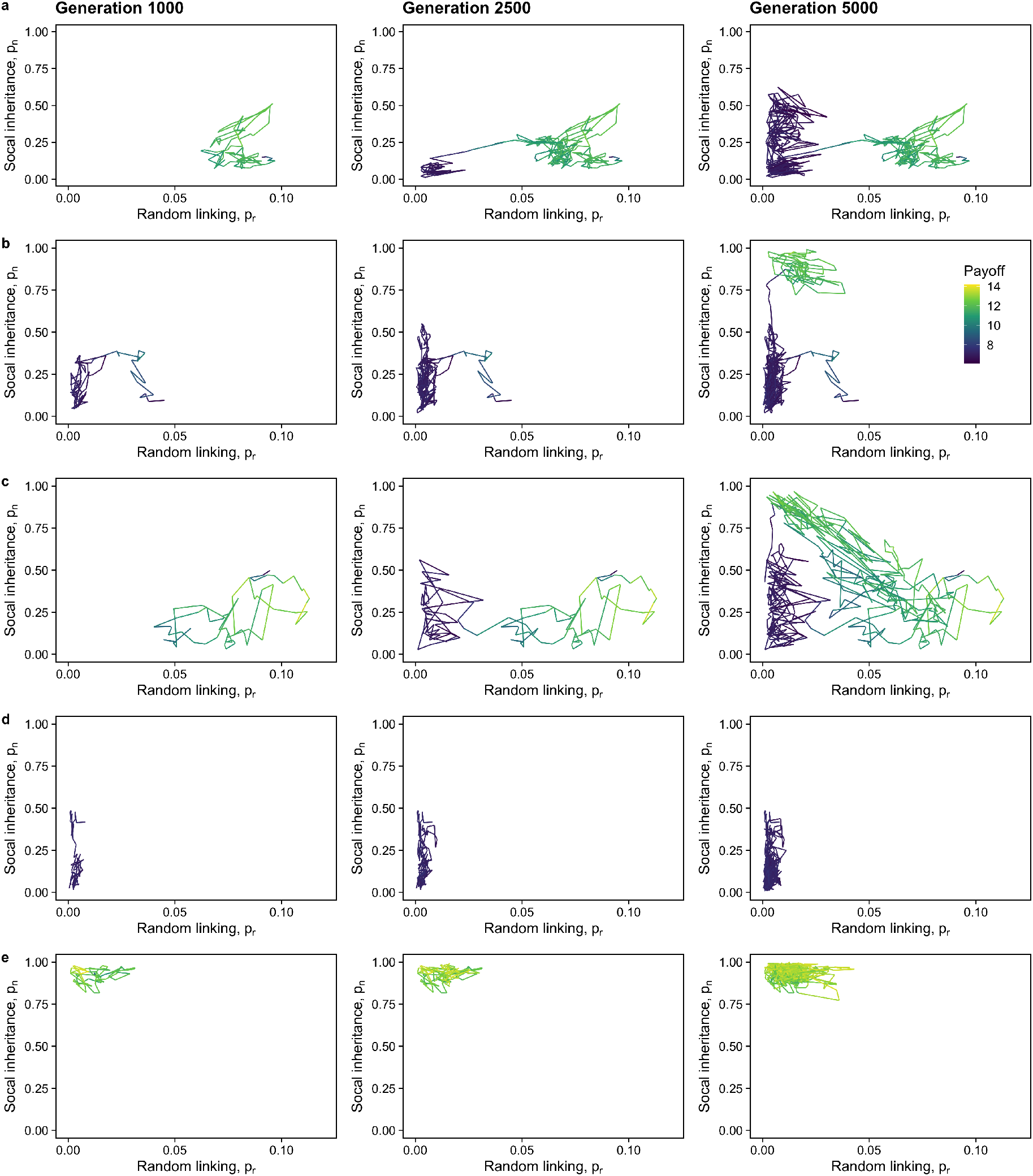
Examples of population connectivity and fitness through time. Data shown are from the same simulations as those in Fig. 2 but for five individual populations (a-e) at three different times throughout the simulation (at 1000, 2500, and 5000 generations). Some populations begin in the high payoff state (high *p*_n_ or *p*_r_) and eventually ‘fall’ into the fitness valley (low *p*_n_, *p*_r_, a-c). Other begin in the low (d) or high (e) payoff state and remain there till the end of the simulation, or even ‘escape’ from the fitness valley (b,c).

**Figure S8.**
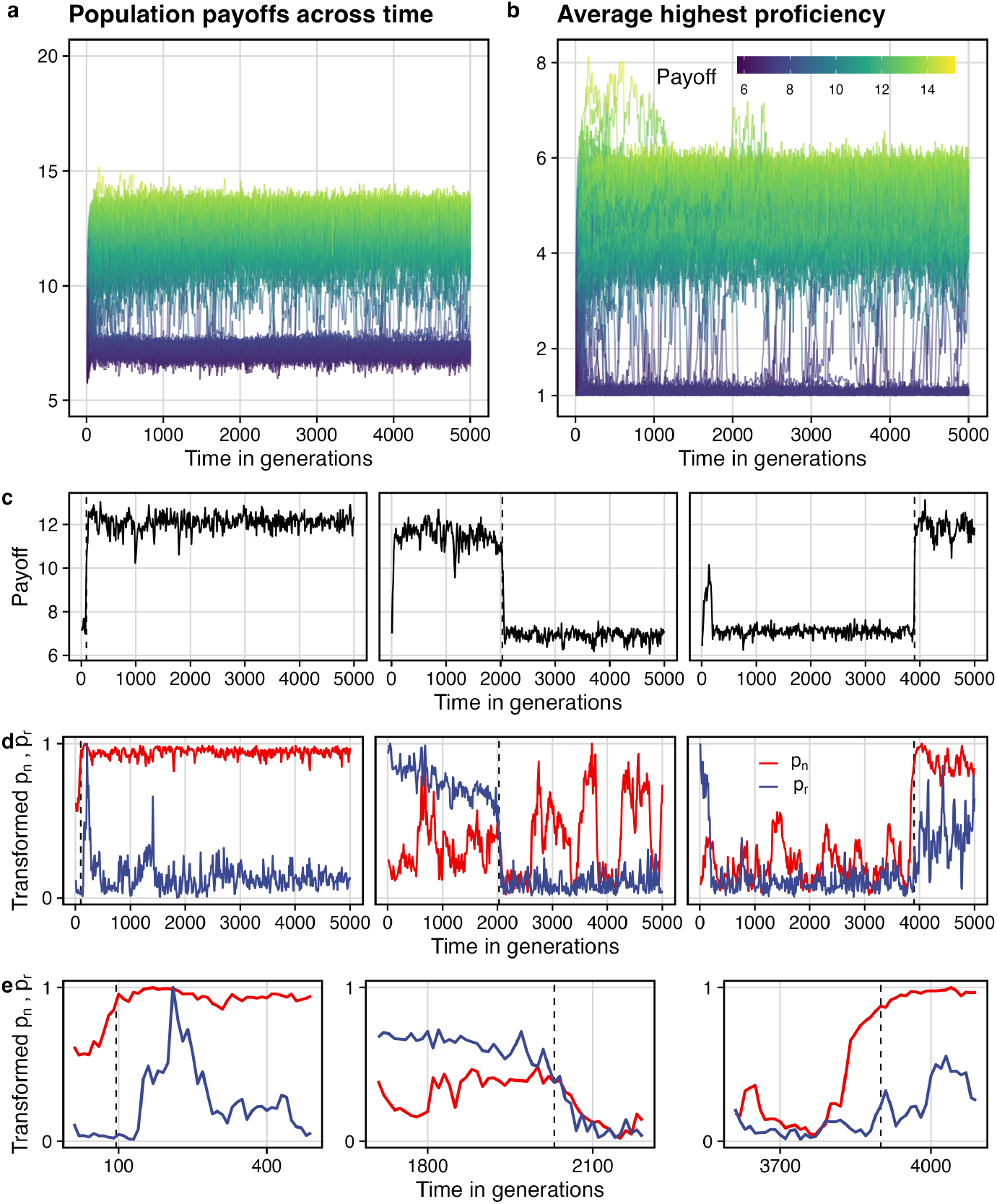
A closer look at transitions between high- and low-payoff states. Panels a and b depict the time-trajectory of an ensemble of simulations that show that they quickly reach a steady state. Panels (c) depicts three individual simulations that provide examples of switches between the low and high-payoff states, with the corresponding transitions in the linking traits (normalised, so that 1 represents the highest observed value) depicted in panels below (d). As can be seen in the close-ups in panels (e), a transition to a high-payoff state begins with an increase in *p*_n_ and a transition to a low payoff state with a decrease in *p*_r_. Simulations with *α* = 0.02 and *β* = 1, *N* = 100, *M* = 500, running for 5,000 generations, *τ* = 0 and *σ* = 0.

**Figure S9.**
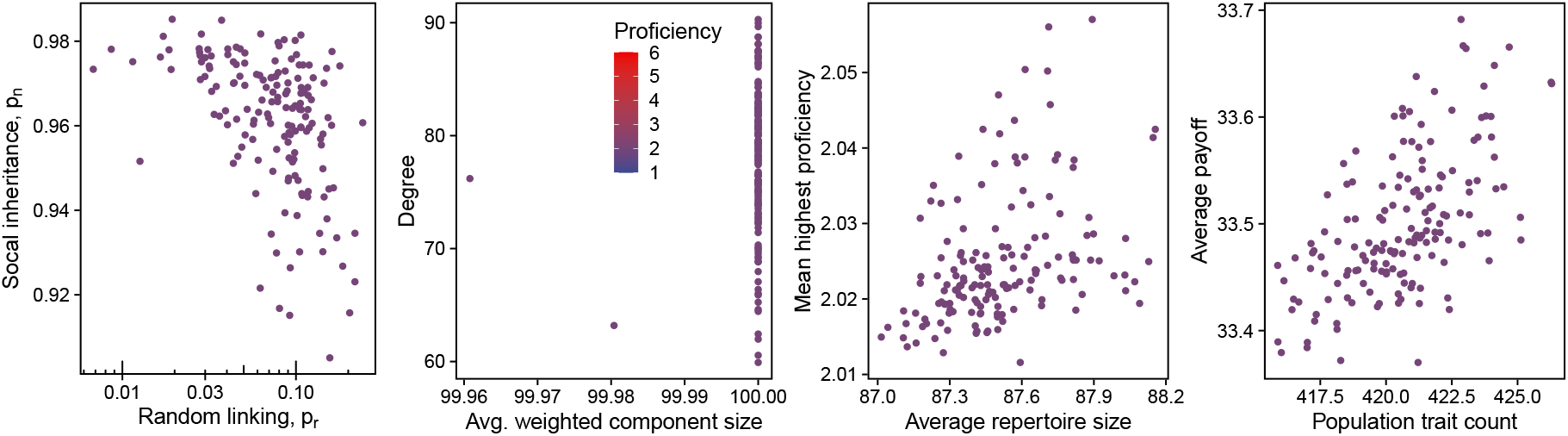
Results from simulations in a homogeneous, constant environment where social learning requires a single exposure. Here, we assume that the probability of socially learning a trait from connections is proportional to its frequency in an individuals neighbourhood (as opposed to proportional to the square of the frequency as in our main results). This corresponds to the assumption that learning can happen with a single exposure. These figures show that the repeated exposure assumption is necessary for the main results we describe in the main text. Specifically, when social learning happens with a single exposure, there is no individual selection for reduced connectivity and all populations are selected to be extremely connected (note that the range on the axes in the first two panels correspond to highly connected networks), have very large repertoire sizes, and display no tradeoff between repertoire size and proficiency (which is generally low, since populations cannot converge on the same traits). Figure S17 shows the equivalent results in heterogeneous and changing environments. In contrast to the social learning described in the main text, here, in a single copying attempt the individual is sampling randomly from the traits in its neighbourhood weighted by the frequency of the trait (*p_i,t_*) and is successfully acquiring the trait with probability *β*.

### S3 Adding environmental heterogeneity to cultural selection on complex graphs

**Figure S10.**
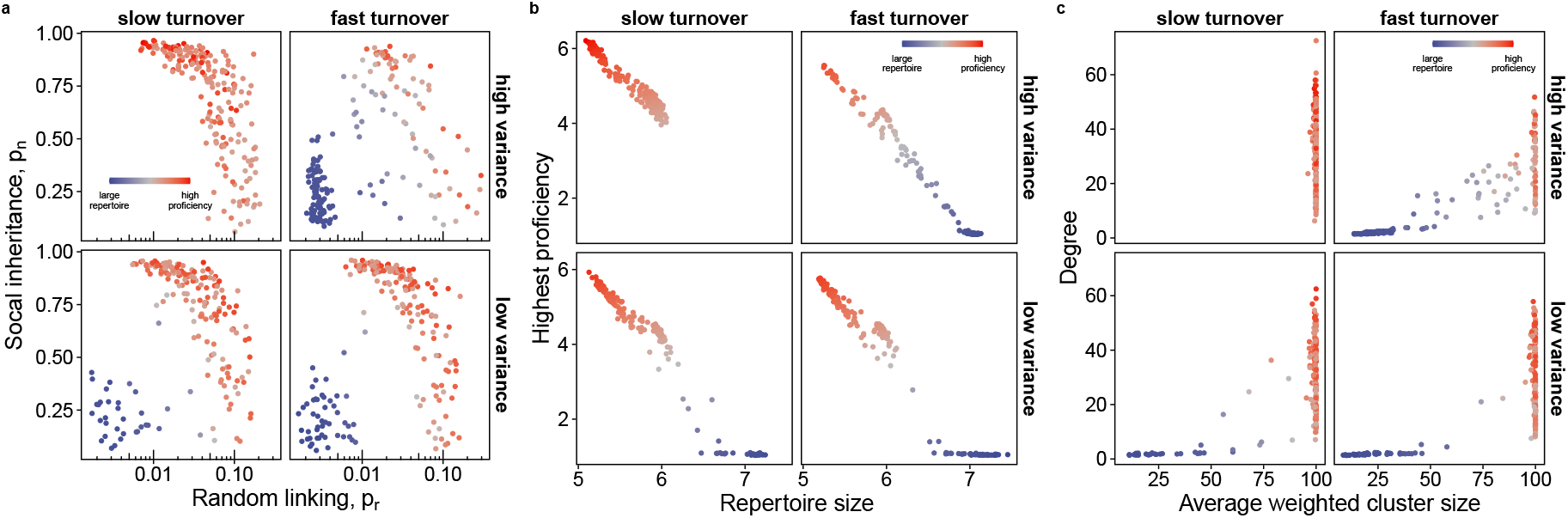
Environmental effects on the evolution of linking parameters. The equivalent of Figure 1 in the main text for four environmental conditions. Colour scale is based on the following data transformation of average repertoire size *R* and highest proficiency *L*: *L*/max(*L*) – *R*/max(*R*). Simulations with *α* = 0.01 and *β* = 1, *N* = 100, *M* = 500, running for 5,000 generations, *τ* ∈ {10^-3^,1} and *σ* = {0.2,1}.

#### S3.1 Varying individual and social learning success probability

Whether either of the strategies is dominant is not only affected by the environment, but also very much by the frequency of innovation and the reliability of social learning.

**Figure S11.**
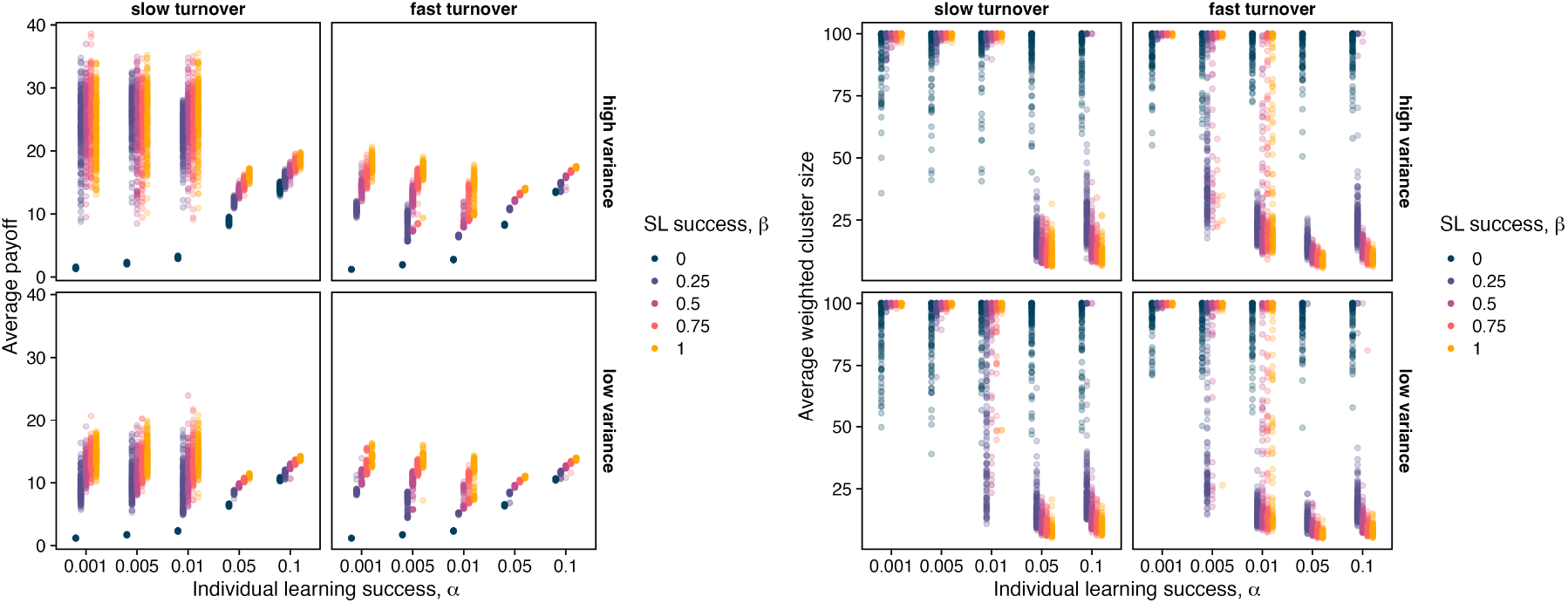
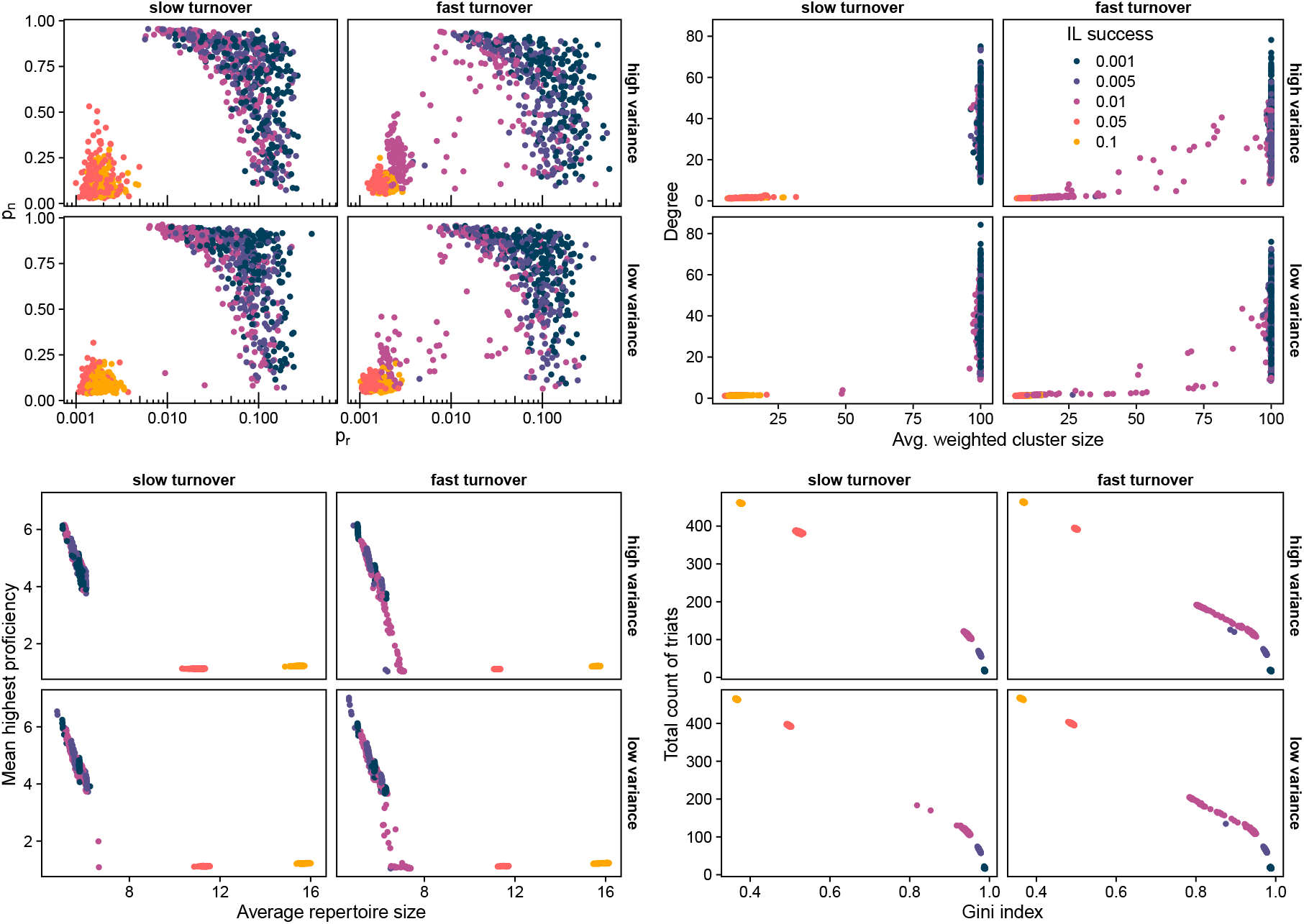
The rate of innovation and social learning affects payoffs in heterogenous environments. The average payoff (a) and average connected component size (b) as a function of social learning rate, for different rates of individual innovation rates. As in the homogenous environment case, in the absence of social learning, we find that increasing innovation always increases payoffs. However, for non-zero social learning success, we find that intermediate values of innovation yield lower payoffs compared to high and low innovation rates. Simulations with *N* = 100, *M* = 500, running for 5,000 generations, *τ* ∈ {10^-3^, 1} and *σ* = {0.4,1}.

#### S3.2 Varying population size

We investigated the effect of population size on coevolution of cumulative culture and social network structure in heterogeneous environments. Figure S12 depicts two of the diagnostic outcomes, average proficiency and connected component size (relative to population size) against population size for different environmental conditions. It shows that only environments with slow turnover produce high proficiency populations, and only when the variance amongst traits is the high proficiency state is dominant. For slow turnover populations, higher population size allows higher proficiency to be built up. However, the proficiency can only go up to a point: with these parameters, population sizes beyond 100 do not cause further increases in the mean proficiency in the high variance slow turnover condition. A more detailed look in figures S13 and S13 corroborate these patterns.

**Figure S12.**
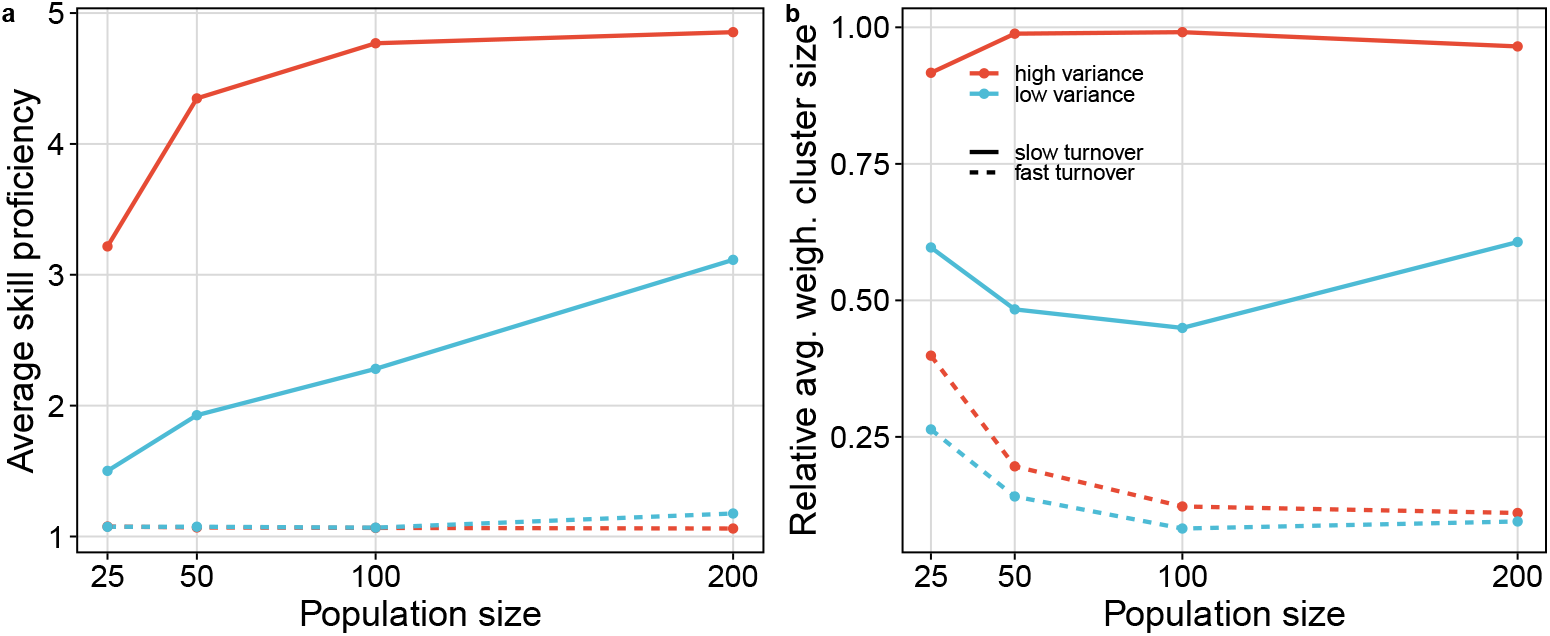
Effect of population size on the coevolution of cumulative culture and social network structure in heterogeneous environments. Each line depicts one prototypical environmental condition: solid lines for slow turnover, dashed for fast; red lines for high variance in payoffs between traits, and blue for low variance. Only slow turnover environments produce high proficiency populations; in these environments, increasing population size allows higher proficiency to be built up. Simulations with *α* = 0.02 and *β* = 1, *N* = 100, *M* = 500, running for 5,000 generations, *τ* ∈ {10^-3^,1} and *σ* = {0.4,1}.

#### S3.3 Fertility selection

In the main text, we assume that the cultural traits affect the probability of the individuals to be selected to die at each time step (mortality selection). An alternative we implemented in our previous work (Smolla and Akçay, 2019) is that the cultural traits affect the probability of being selected to reproduce. To check whether implementing selection at the mortality stage vs. reproduction affects our main results, we ran our simulations with selection at the reproduction stage. Specifically, in each time step, one individual is selected at random to die, but the individual to reproduce is selected with probability proportional to *M* (*W_i_*) as defined in the methods. Figure S15 shows the equivalent of Figure 1 in the main text. It shows that our results are not affected by mortality vs. fertility selection. Specifically, we again observe the same two distinct population states (low connectivity, broad repertoire, low proficiency and payoff vs. high connectivity, narrower repertoire, high proficiency and payoff). Moreover, we also see the cycling between these states, driven by the exact same patterns in payoffs depicted in Figures 2 and 3.

**Figure S13.**
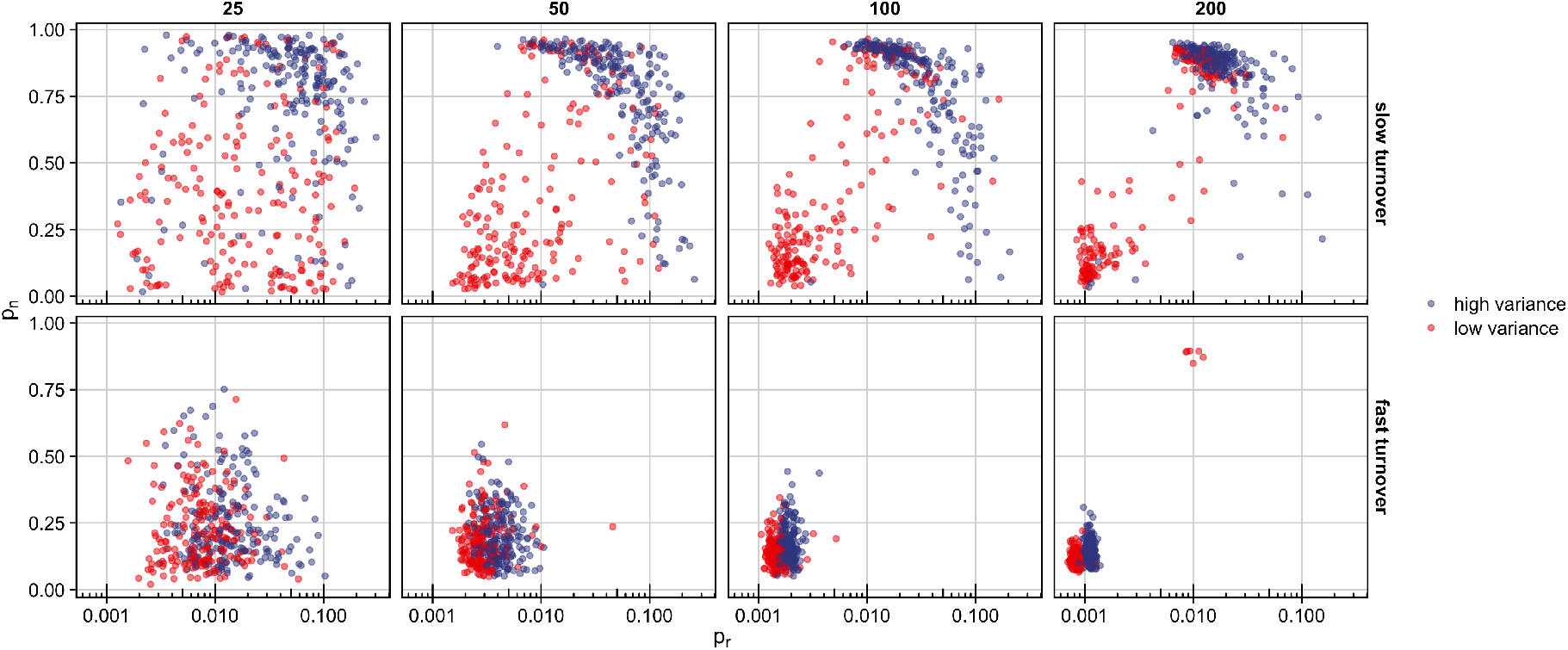
More detailed look at the populations’ linking traits at different population sizes across environments. In slow turnover environments, increasing population size tends to favour more connected populations, whereas in fast turnover environments, increasing population size has a much weaker effect. For slow turnover, population size effect is stronger with high variance between traits, while for fast turnover, the population size effect is only apparent with low variance. Simulations with *α* = 0.02 and *β* = 1, *N* = 100, *M* = 500, running for 5,000 generations, *τ* ∈ {10^-3^, 1} and *σ* = {0.4,1}.

#### S3.4 No saturating payoffs

As described in the methods, we assume that the mortality probabilities are functions of not directly the payoff *W_i_*, but a saturating function of it *M*(*W_i_*), which we assume has a Michealis-Menten form. We assume a relatively large half-rate constant (*K* = 50) which means that for most parameter regimes, *M*(*W_i_*) is closer to linear in *W_i_*, as the latter is not close to the half-rate constant. As explained in the methods, we made this assumption to guard against the possibility that in heterogeneous environments with large payoff variance between traits, a single “lucky” innovation with very high results might introduce unrealistic amount of strong selection. To check whether this assumption is consequential for our base results, we ran our simulations in homogeneous environments assuming selection (probability of mortality in a given time step) is directly proportional (to the inverse of) *W_i_*, instead of a saturating function of it. Figure S16 depicts the equivalent of Figure 1 in the main text, and shows that this assumption is not consequential for our main result.

#### S3.5 Simple contagion based social learning in heterogeneous environments

Figure S17 shows the equivalent of S9 in different heterogeneous and changing environments. These simulations again assume that social learning probability is proportional to the frequency of the trait in the neighbourhood rather than the square of the frequency. It shows that all of the results in homogeneous environments carry over to heterogeneous environments, with slight changes as expected due to the payoff variation between traits and turnover rate. Specifically, we again do not find the conflict between group payoff and individual fitness, and therefore no cycling.

**Figure S14.**
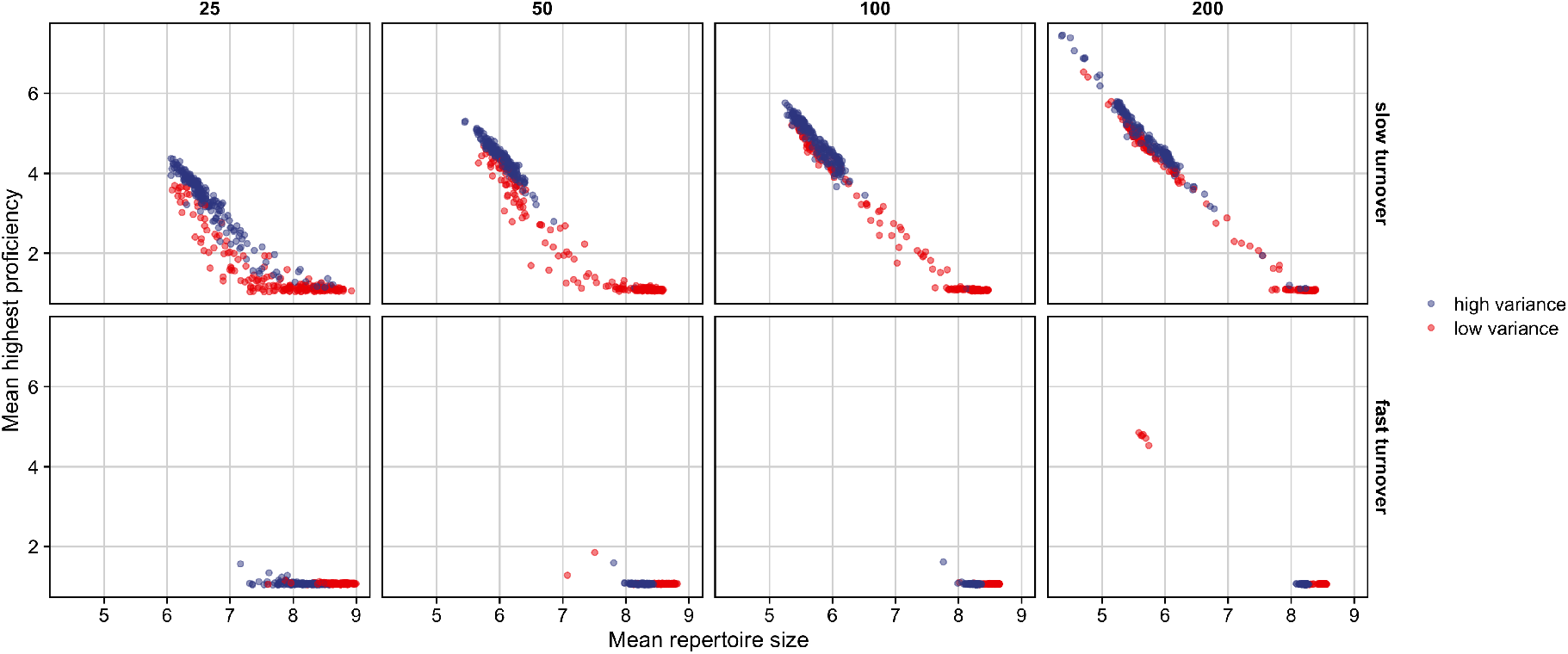
Proficiency and repertoire size with population size across environments. The mean proficiency and repertoire size for the same populations as in Figure 2. Increasing population size in slow changing environments favours increased proficiency (especially with high variance), which in fast changing environments it has relatively little effect, and that only under low variance. Simulations with *α* = 0.02 and *β* = 1, *N* = 100, *M* = 500, running for 5,000 generations, *τ* ∈ {10^-3^, 1} and *σ* = {0.4,1}.

**Figure S15.**
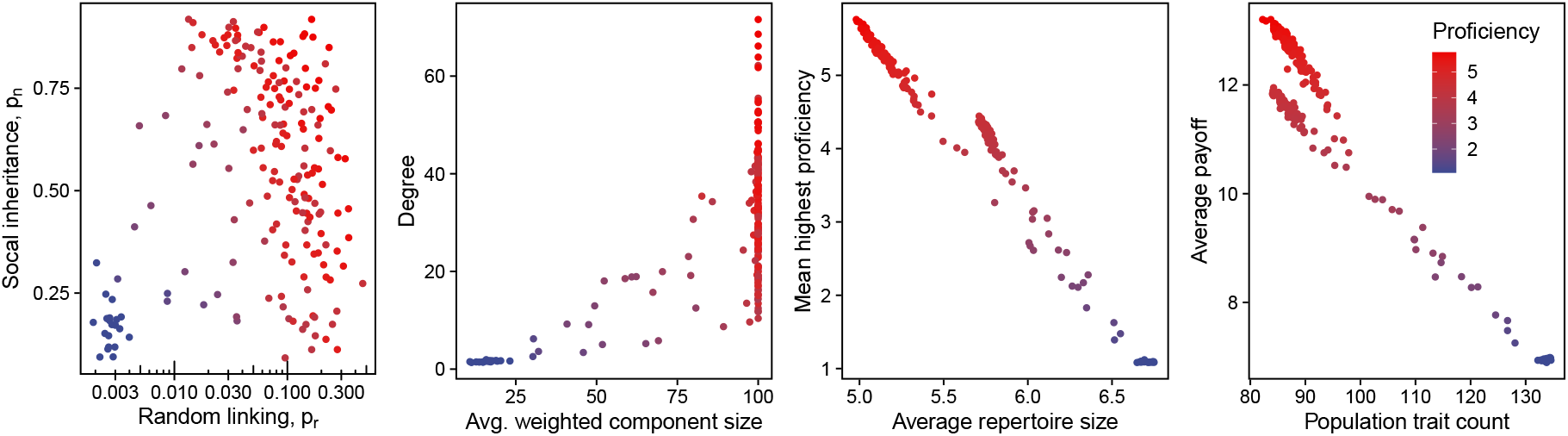
Results with fertility selection in homogeneous environments. The equivalent of Figure 1 in the main text, but with selection due to cultural traits on fertility instead of mortality. Specifically, in these simulations, mortality is random but probability is getting selected to reproduce is proportional to *M*(*W_i_*) as described in the methods. All our results in homogeneous environments carry over to fertility selection, including the two pathways to cultural adaptation and cycling between them. Simulations with *α* = 0.01 and *β* = 1, *N* = 100, *M* = 500, running for 10,000 generations, *τ* = 0 and *σ* = 0, 200 replicates.

**Figure S16.**
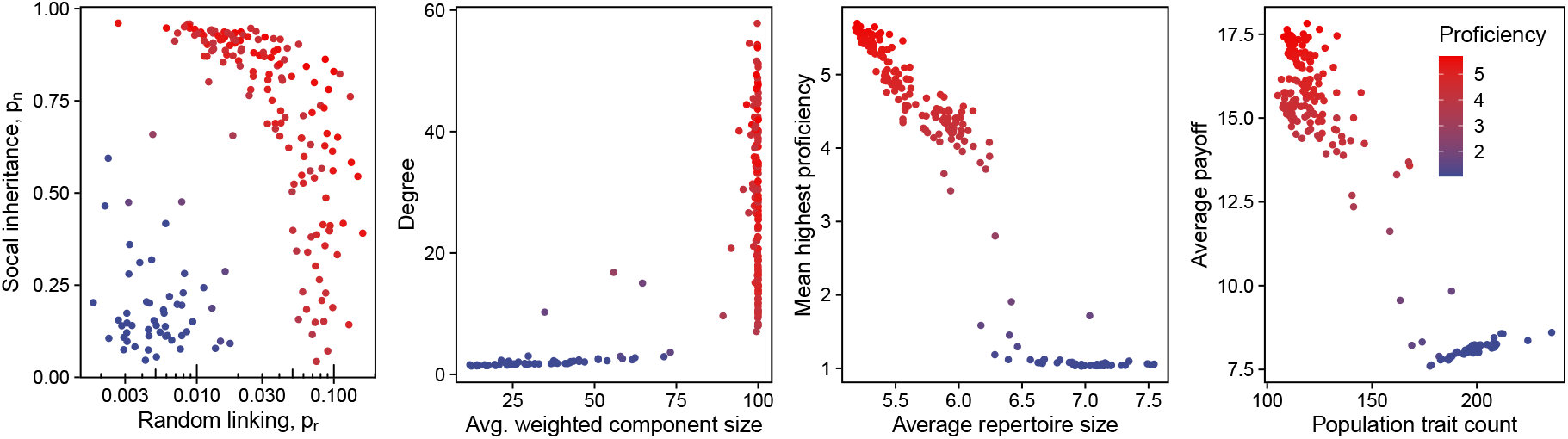
Results without saturating fitness values. The equivalent of Figure 1 in the main text, but with fitness simply equal to payoff *W_i_*, i.e., without the saturating Michealis-Menten function. All our results from the main text again carry over to the case without the saturating fitness function. Simulations with *α* = 0.01 and *β* = 1, *N* = 100, *M* = 500, running for 5,000 generations, *τ* = 0 and *σ* = 0, 200 repetitions.

**Figure S17.**
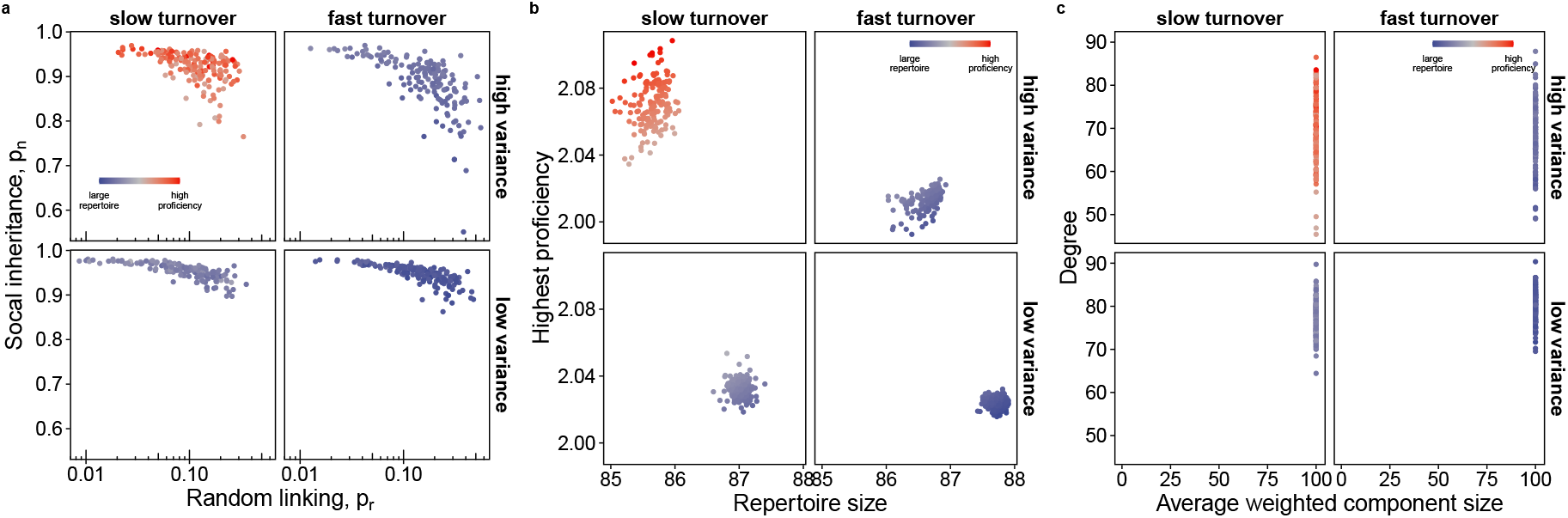
Results from simulations where social learning requires only a single exposure. Compared to the results from the main text (where we assume social learning requires multiple exposures), we do not find the two different pathways (dense networks with high proficiency and small repertoire vs. sparse networks with large repertoires and low proficiency). Therefore, we also do not find the cycling behaviour that we describe in the main text results. Instead, we find that dense network evolve (across all tested environments) and that proficiency remains low but repertoires are substantially larger. For simple contagion the probability to acquire a specific trait is *βp_i,t_*, whereas for complex contagion it is 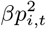. This makes acquiring traits with single exposure much more likely, which leads to the larger repertoires. On the other hand, having a large repertoire and a limited number of learning turns limits the probability to successfully ‘level up’ in any trait, which causes proficiency to remain low. Colour scale is based on the following data transformation of average repertoire size *R* and highest proficiency *L*: *L*/max(*L*) – *R*/max(*R*). Simulations with *α* = 0.01 and *β* = 1, *N* = 100, *M* = 500, running for 5,000 generations, *τ* ∈ {10^-3^, 1} and *σ* = {0.2,1}, each with 160 repetitions.

